# Altered Hepatic Metabolism in Down Syndrome

**DOI:** 10.1101/2025.05.27.656393

**Authors:** Lauren N. Dunn, Brian F. Niemeyer, Neetha P. Eduthan, Kyndal A. Schade, Katherine A. Waugh, Chrisstopher Brown, Angela L. Rachubinski, Ariel E. Timkovich, David J. Orlicky, Matthew D. Galbraith, Joaquin M. Espinosa, Kelly D. Sullivan

**Affiliations:** Linda Crnic Institute for Down Syndrome, University of Colorado Anschutz Medical Campus, Aurora, Colorado, USA; Department of Cell Biology and Physiology, University of Kansas Medical Center, Kansas City, KS, USA; Department of Pediatrics, Section of Developmental Pediatrics, University of Colorado Anschutz Medical Campus, Aurora, Colorado, USA; Department of Pathology, University of Colorado Anschutz Medical Campus, Aurora, Colorado, USA; Department of Pharmacology, University of Colorado Anschutz Medical Campus, Aurora, Colorado, USA; RNA Bioscience Initiative, University of Colorado Anschutz Medical Campus, Aurora, Colorado, USA; Department of Pediatrics, Section of Developmental Biology, University of Colorado Anschutz Medical Campus, Aurora, Colorado, USA

## Abstract

Trisomy 21 (T21) gives rise to Down syndrome (DS), the most commonly occurring chromosomal abnormality in humans. T21 affects nearly every organ and tissue system in the body, predisposing individuals with DS to congenital heart defects, autoimmunity, and Alzheimer’s disease, among other co-occurring conditions. Here, using multi-omic analysis of plasma from more than 400 people, we report broad metabolic changes in the population with DS typified by increased bile acid levels and protein signatures of liver dysfunction. In a mouse model of DS, we demonstrate conservation of perturbed bile acid metabolism accompanied by liver pathology. Bulk RNA-sequencing revealed widespread impacts of the Dp16 model on hepatic metabolism and inflammation, while single-cell transcriptomics highlighted cell types associated with these observations. Modulation of dietary fat profoundly impacted gene expression, bile acids, and liver pathology. Overall, these data represent evidence for altered hepatic metabolism in DS that could be modulated by diet.

## INTRODUCTION

Triplication of chromosome 21 (chr21), known as trisomy 21 (T21), results in the genetic disorder commonly known as Down syndrome (DS).^1^ T21 is the most common chromosomal abnormality in the human population, currently affecting approximately 300,000 individuals in the United States and around 6 million worldwide.^2^ It has been observed that individuals with T21 experience a distinctive disease spectrum, with perturbations seen in essentially every cell type, tissue, and organ system in the body.^3^ Although individuals with DS endure a predisposition to many co-occurring conditions, including obesity, Alzheimer’s disease, autoimmune disorders, congenital heart defects, and autism, the mechanisms driving the increased rates of these phenotypes, or the phenotypic diversity, in the population with DS remain poorly understood.^3^ Recent advances in high-throughput ‘omics’ approaches have provided new insights into the system-wide impacts of T21 on molecular and cellular biosignatures. Our group has demonstrated widespread effects of T21 on the transcriptome,^4–6^ proteome,^4,6–9^ metabolome,^6,8^ and peripheral immune compartment,^5,6,10^ leading to the observations that DS is 1) an interferonopathy,^4,6^ 2) characterized by IGF1 deficiency,^9^ and 3) comprised of distinct molecular subtypes.^11^ A core component of these observations is global alterations in metabolism, including specific changes to the kynurenine pathway^6,8^ and gene expression changes associated with heme metabolism.^12^ These studies and others have demonstrated widespread metabolic dysregulation in DS, though many questions remain with respect to causes and consequences.^12–17^

Bile acids (BAs) have diverse roles in metabolism, homeostasis, and disease.^18^ Primary BAs are synthesized from cholesterol in hepatocytes and secreted into bile ducts where they are conjugated to taurine or glycine, which increases their solubility and decreases their cell membrane permeability.^19^ These BAs are then trafficked to the gut where they play central roles in the metabolism of lipids, fats, and cholesterol. In the gut, the microbiome can modify BAs into so-called secondary BAs. After their trafficking functions are complete, BAs are deconjugated and up to 95% are returned to the liver via enterohepatic circulation.^19^ In the liver and intestine, BAs serve as ligands for nuclear receptors, such as Farnesoid X Receptor (FXR),^20^ where they stimulate gene expression programs that regulate further production of BAs and maintain lipid homeostasis. BAs are also ligands for G Protein-Coupled Receptors (GPCRs), such as TGR5,^21^ which activates phosphatidylinositol 3-kinase (PI3K)/AKT signaling in enterocytes.

Alterations in BA metabolism, observed in the form of elevated plasma BA levels, have long been considered a hallmark of altered liver metabolism.^18,22^ Although around 4.5 million adults in the US (1.8%) have a diagnosed liver disease, the American Liver Foundation estimates that another 80-100 million adults in the U.S. have an undiagnosed liver disease.^23^ The most common forms of liver disease include non-alcoholic fatty liver disease (NAFLD), now referred to as metabolic dysfunction-associated steatotic liver disease (MASLD), which is comprised of a spectrum of liver diseases that can range from simple fatty liver (steatosis) to more severe steatohepatitis (MASH) that can lead to liver scarring in the form of fibrosis and cirrhosis.^24^ The prevalence of MASLD is increasing globally, particularly in Western nations.^25^

The rates of liver disease in individuals with DS remains unknown though. Liver disease diagnoses are commonly made on the results of Liver Function Tests (LFTs), such as elevated alanine aminotransferase (ALT) or aspartate aminotransferase (AST), which are markers of hepatocellular damage commonly seen in patients with MASLD, or elevated plasma BAs, which are commonly associated with cholestasis.^26,27^ Imaging-based methods such as ultrasound or FibroScan assist in proper diagnosis of liver disease.^27,28^ A recent large cohort study of individuals with DS examined LFTs, including ALT and AST, and concluded that T21 has no biochemical evidence of hepatocellular injury,^17^ while another study using electronic medical records concluded that individuals with DS are less likely to suffer from liver disease.^29^ In contrast, a 2017 study from Italy demonstrated that more than 60% of DS individuals in a cohort ranging in age from 5-18 years had ultrasound findings consistent with fatty liver.^30^ Strikingly, this fatty liver phenomenon was observed even in the absence of obesity and was not associated with abnormal AST and ALT levels. These conflicting reports suggest the presence of liver dysfunction in the population with DS that is not easily detectable through conventional blood-based LFTs. Further research is therefore needed to define the prevalence and consequences of liver dysfunction in this population, as well as provide evidence as to which common features are dysregulated.

In this study, we examined plasma metabolomic and proteomic data from a large cohort of individuals with DS as well as euploid controls. We report that the population with DS has increased plasma BAs. Within the plasma proteome, we observed evidence of broad liver dysfunction among the most significantly dysregulated pathways. We have previously shown that adult Dp(16)1Yey/+ mice (referred to hereinafter as Dp16), a mouse model of DS, displays a liver phenotype characterized primarily by increased inflammation, arterial thickening, and ductular reaction.^31^ Here, we expand upon this phenotype in a new cohort of adult Dp16 mice. Importantly, metabolomic analysis of plasma from Dp16 mice shows elevated levels of plasma BAs. Transcriptome analysis of the Dp16 liver identified signatures of both extracellular matrix and immune system dysregulation, as well as metabolic dysfunction and fibrosis among the top dysregulated pathways. Single-cell RNA sequencing (scRNAseq) of Dp16 liver samples confirmed these findings and highlighted cell type-specific alterations in metabolic and inflammatory pathways. Human hepatocytes differentiated from a panel of T21 and euploid control induced pluripotent stem cells (iPSCs) demonstrated concordance with the features dysregulated in the plasma of individuals with DS, including protein abundance, lipid production, and BA production. We also found commonalities in gene expression profiles within two hepatocyte clusters from the Dp16 liver scRNAseq data, suggesting that some aspects of liver dysfunction in DS are cell-type specific.

Finally, we challenged Dp16 mice with a high-fat diet (HFD) and found that the observed liver phenotype is driven primarily by diet. Transcriptome analysis of the liver after HFD challenge showed perturbations in pathways associated with the inflammation and metabolism. Importantly, we also show that plasma BAs are perturbed after this diet, suggesting a correlation between diet and BA dysregulation. Overall, the evidence presented here indicates that individuals with DS, as well as the Dp16 mouse model, experience widespread liver dysfunction, defined primarily by dysregulated hepatic metabolism, particularly BA metabolism, and this dysregulation may be driven largely by dietary fat.

## RESULTS

### Altered bile acid metabolism in Down syndrome

To expand upon our previous observations that DS globally alters metabolism, we reanalyzed datasets generated by the Human Trisome Project (HTP).^11,32–35^ These datasets represent molecular profiling of tens of thousands of features across multiple -omics platforms from 316 individuals with DS (T21) and 103 euploid controls (D21). We first examined the impact of T21 on the plasma metabolome by analyzing the relative levels of plasma metabolites detected via ultra-high performance liquid chromatography (UHPLC)-Mass Spectrometry. Differentially abundant metabolites in the T21 population were determined using a linear model with adjustment for age, sex, and body mass index (BMI) as co-variates. We identified 102 differentially abundant metabolites (Benjamini-Hochberg correction with a 10% FDR) (**Figure 1A, Table S1A**). Next, we grouped metabolites based on pathway, which highlighted global metabolic changes in DS, typified by elevation of fatty acids/eicosanoids, carnitines and fatty acids, and BAs, as well as depletion of amino acids and nucleotides (**Figure 1B**). The BAs were the largest group of related, significantly different metabolites dysregulated in the same direction. Closer examination of BAs revealed a total of 11 measured, with nine elevated in those with DS, including taurolithocholic acid (**Figure 1C, 1D, and S1A**). We observed a trend towards an increase in the total abundance of primary BAs in the plasma of individuals with T21, and a significant increase in secondary BAs (**Figure S1B**). Notably, these elevated levels were conserved across the lifespan of individuals with DS, with no real impact of age on the levels of taurolithocholic acid, for example (**Figure 1E, Table S1B)**. In the typical population, elevated BMI is frequently associated with liver disease, however no significant association between BMI and taurolithocholic acid was observed in our T21 cohort (**Figure S1C**).

**Figure 1.**
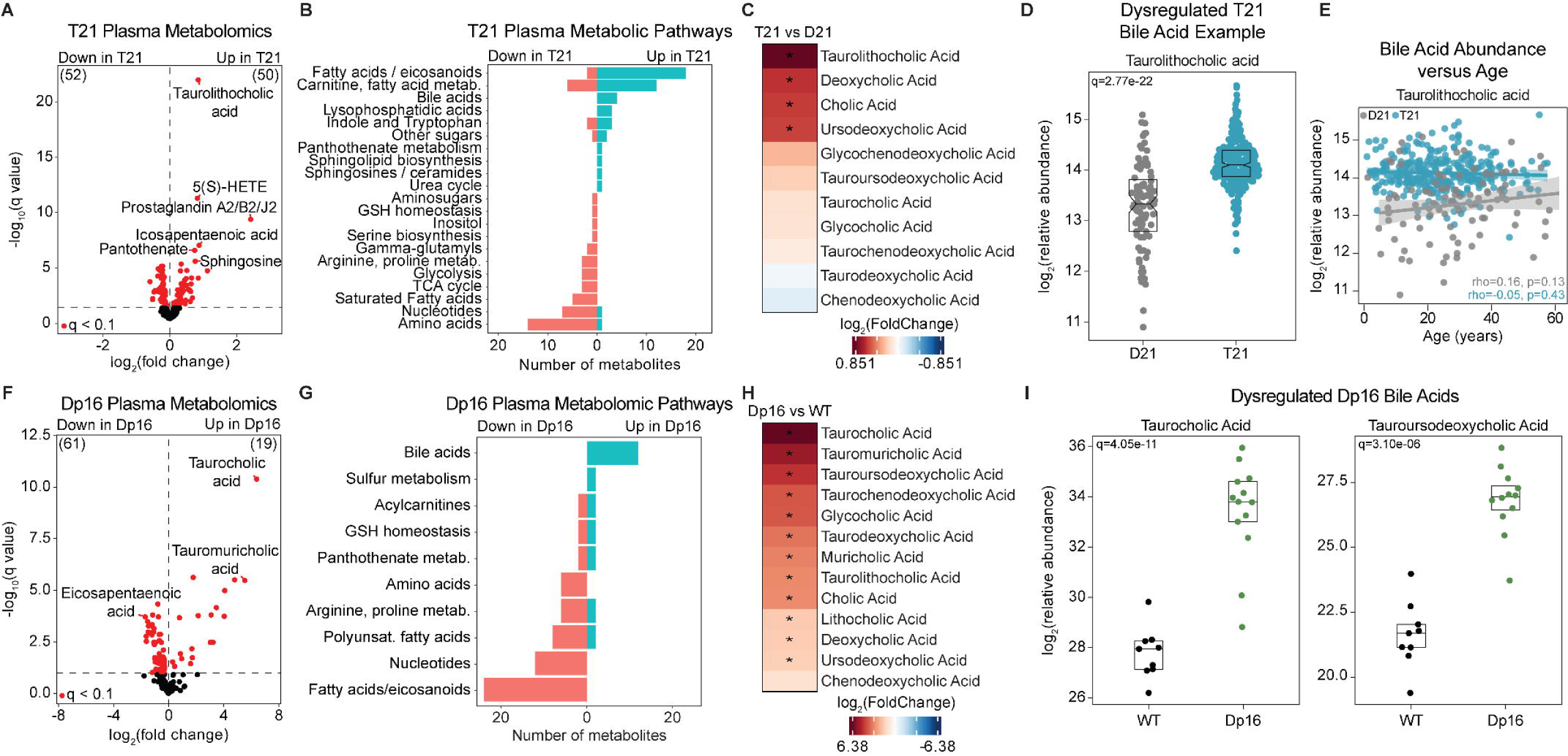
Altered bile acid metabolism in Down syndrome. (A) Volcano plot summarizing results of plasma metabolomic analysis of individuals with DS (n=316) versus disomic individuals (n=103). Significantly differentially abundant (q<0.1) metabolites are colored in red. (B) Waterfall plot summarizing metabolites with significant differences in relative abundance in plasma of individuals with T21 versus D21 controls (q<0.1). Bars are color-coded by increased relative abundance in DS (blue) versus decreased in DS (red). (C) Heatmap displaying log_2_(FoldChange) of plasma bile acids in those with T21 relative to D21 individuals. Color coding indicates a positive (red) or a negative (blue) log_2_(FoldChange) value. Asterisks denote significance (q<0.1). (D) Sina plot displaying log_2_(relative abundance) of taurolithocholic acid in the plasma of D21 individuals and those with DS. Boxes represent interquartile ranges and medians, with notches approximating 95% confidence intervals. Benjamini-Hochberg adjusted p-values (q-values) are indicated. (E) Scatterplot displaying the relationship between relative abundance of taurolithocholic acid and age in years both for D21 individuals and those with DS. Spearman rho values and p-values are shown. (F) Volcano plot summarizing results of plasma metabolomic analysis of Dp16 mice (n=13) compared to wildtype (n=9). Significantly differentially abundant (q<0.1) metabolites are colored in red. (G) Waterfall plot summarizing metabolites with significant differences in relative abundance in plasma from Dp16 mice versus wildtype. (H) Heatmap displaying log_2_(FoldChange) of plasma bile acids in Dp16 mice relative to wildtype. Color coding indicates a positive (red) or a negative (blue) log_2_(FoldChange) value. Asterisks denote significance (q<0.1). (I) Sina plots displaying log_2_(relative abundance) of taurocholic acid and tauroursodeoxycholic acid plasma metabolites in wildtype and Dp16 mice. Benjamini-Hochberg adjusted p-values (q-values) are indicated.

To investigate potential mechanisms leading to BA dysregulation in individuals with T21, we looked to the Dp16 mouse model of DS. This model harbors a segmental duplication of a portion of mouse chromosome 16, conferring trisomy of around 120 protein coding genes orthologous to human chromosome 21.^36^ We first characterized the plasma metabolome of a cohort of Dp16 animals and matched wildtype (WT) littermate controls using UHPLC-Mass Spectrometry to measure the relative abundance of around 150 metabolites. Using a linear model with adjustment for age and sex as co-variates, we identified 80 differentially abundant metabolites (Benjamini-Hochberg correction with a 10% FDR) (**Figure 1F, Table S1C**). Strikingly, BAs were the largest pathway elevated in Dp16 animals, with 12 of 17 significantly elevated (**Figures 1G and 1H**). We observed differential abundance of both primary BAs, cholic and taurocholic acid, and secondary BAs, including taurolithocholic acid, deoxycholic acid, tauroursodeoxycholic acid, and the murine-specific muricholic acid (**Figure 1G 1I, S1C, and S1D**). We also observed a significant increase in both primary and secondary BAs in the Dp16 mice (**Figure S1E**). Thus, the Dp16 mouse model of DS exhibits defects in BA metabolism similar to those seen in the population with T21.

### Evidence for widespread liver dysfunction in Down syndrome

Given that high levels of plasma BAs can indicate the presence of liver disease, we next reanalyzed our HTP SOMAscan proteomic^37^ data for markers of liver function. We measured more than 4,500 epitopes in the plasma of 419 samples from the HTP and, using a linear model with adjustment for age, sex, and BMI as co-variates, found hundreds of proteins with altered abundance in individuals with T21, including nearly 1,000 that are more abundant (Benjamini-Hochberg correction with a 10% FDR) (**Figure S2A, Table S2A**). Ingenuity Pathway Analysis (IPA) of these differentially abundant proteins uncovered several signatures of liver dysfunction in individuals with DS (**Figure 2A**). Indeed, two of the top six enriched canonical pathways were related to liver dysfunction: Liver X Receptor/Retinoid X Receptor (LXR/RXR) Activation and Hepatic Fibrosis/Stellate Cell Activation (**Figure 2A**). LXRs, nuclear receptors that can form heterodimers with RXRs, play a key role in lipid metabolism. Upon activation of LXRs by elevated levels of intracellular cholesterol, cholesterol absorption, transport, and excretion occurs. Dysregulation of the coagulation system can also indicate liver dysfunction, as hepatocytes and liver sinusoidal epithelial cells, two of the main cell types in the liver, play a key role in synthesizing coagulation proteins. Dysregulation of the hepatic fibrosis pathway suggests there may be a buildup of collagen protein that has resulted in hepatic fibrosis. Gene set enrichment analysis (GSEA) of this dataset revealed similar findings: positively enriched hallmarks included multiple immune-related pathways, and negatively enriched pathways included the coagulation pathway (**Figure S2B, Table S2B**).

**Figure 2.**
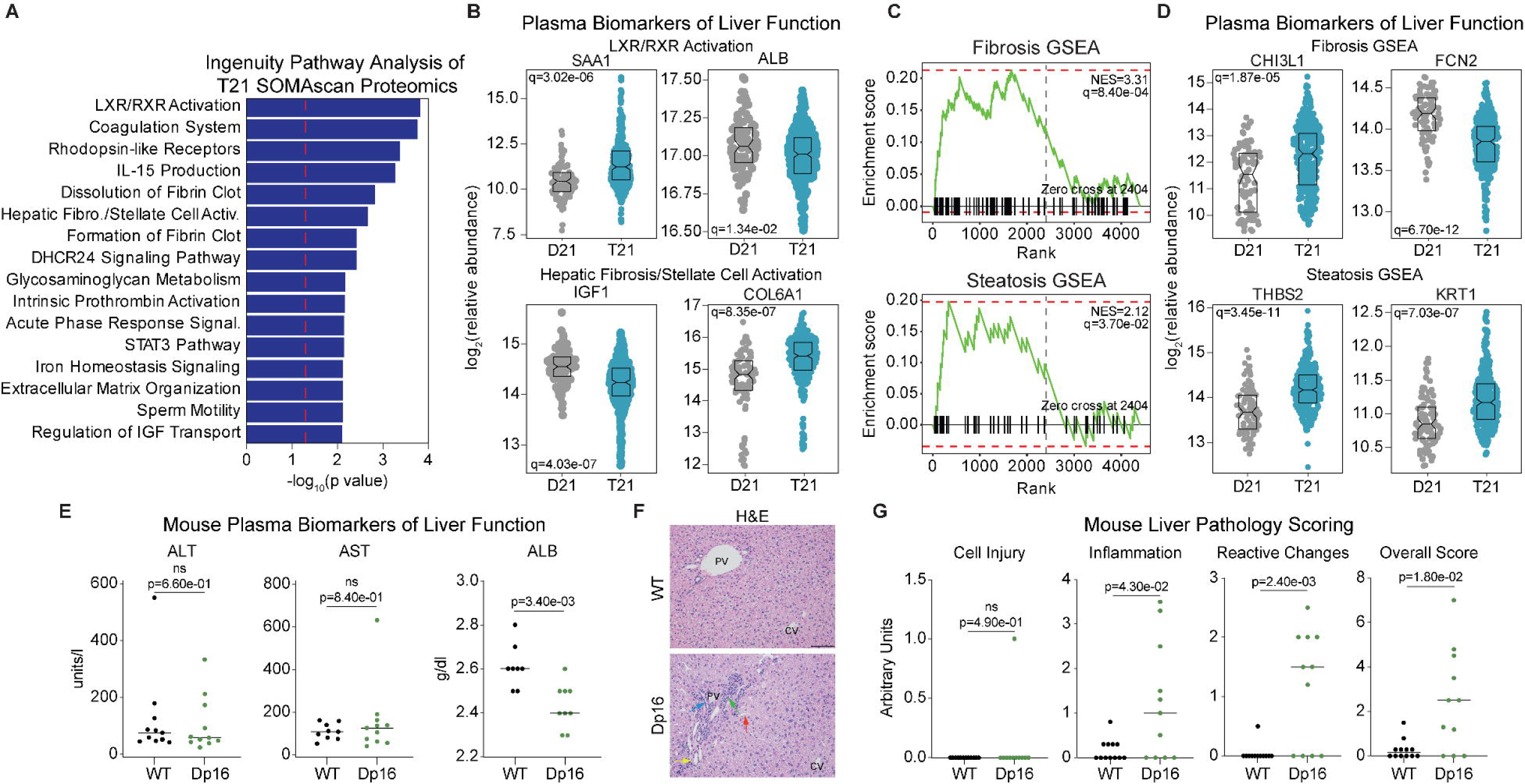
Evidence for widespread liver dysfunction in Down syndrome. (A) Bar plot summarizing Ingenuity Pathway Analysis of plasma SOMAscan proteomics from individuals with DS (n=316) versus D21 individuals (n=103). Dotted line indicates the significance threshold of p < 0.05. (B) Sina plots displaying the relative abundance of various plasma proteins in disomic individuals and those with DS. Benjamini-Hochberg adjusted p-values (q-values) are indicated. (C) Enrichment plots showing enrichment scores for Gene Set Enrichment Analysis signatures of Fibrosis and Steatosis in plasma proteomics of individuals with DS. NES – Normalized Enrichment Score (D) Sina plots displaying the relative abundance of various plasma proteins in D21 individuals and those with DS. Benjamini-Hochberg adjusted p-values (q-values) are indicated. (E) Sina plots showing the concentration of various biomarkers of liver function in wildtype (n=8-11, 3-6 females) and Dp16 (n=9-11, 3-6 females) plasma. Individual data are presented with a bar at the median and p-values, as determined by Mann-Whitney U test, are shown. (F) Representative images of H&E-stained liver sections from wildtype and Dp16 mice. Arrows indicate the following features: red, enlarged periportal sinusoids; blue, bile duct; yellow, hepatic artery branches; green, periportal inflammatory cells. PV indicates portal vein and CV indicates central vein. Scale bar is 100 microns. (G) Sina plots displaying metrics of liver pathology scoring from adult wildtype (n=12, 6 females) and Dp16 (n=11, 5 females) mice. Individual data are presented with a bar at the median and p-values, as determined by Mann-Whitney U test, are shown. Boxes in Sina plots represent interquartile ranges and medians, with notches approximating 95% confidence intervals.

Consistent with prior reports,^34,38^ we found no significant difference in the plasma levels of liver enzymes alanine transaminase (ALT) and aspartate aminotransferase (AST) in the T21 group (**Figure S2C**). However, we did observe significant differences in several proteins known to be dysregulated in liver disease, some of which have the potential to predict the severity of various disorders^39,40^. For example, we observed increased levels of serum amyloid A1 (SAA1) and collagen 6A1 (COL6A1), and decreased levels of albumin (ALB) and insulin-like growth factor 1 (IGF1) (**Figure 2B**). SAA1 is an acute phase protein primarily synthesized in the liver and is released into the bloodstream upon damage or infection and increased levels of COL6A1in the plasma have been seen to be associated with more severe liver disease.^41^ Hypoalbuminemia is a hallmark of liver dysfunction, as albumin is produced by the liver. Lower levels of IGF1 have been associated with the progression of various liver diseases, as 75% of IGF1 is produced by the liver and serves a protective role within the organ.^42,43^

Next, we took advantage of a study that clinically staged both steatosis and fibrosis within euploid individuals diagnosed with liver disease and generated plasma proteomic biosignatures of disease using the SOMAscan platform.^33,44^ The aptamer-based nature of this assay allows us to easily compare results across studies. Therefore, using the lists of differentially abundant proteins identified by Govaere *et al*, we performed Gene Set Enrichment Analysis (GSEA) on our HTP SOMAscan data. We found that Fibrosis and Steatosis signatures were significantly enriched in individuals with DS (**Figure 2C, Table S2C**). Within these signatures, we observed proteins involved in a wide range of biological functions linked to liver dysfunction, including innate immunity/extracellular matrix remodeling (CHI3L1, FCN2, and THBS2), cell adhesion (SELE), and metabolism (AMY2A) (**Figures 2D and S2C**).

We also measured canonical plasma markers of liver function in Dp16 animals and found that similarly to people with DS, Dp16 animals display no significant differences in ALT nor AST compared to their WT littermates (**Figure 2E, Table S2D**). This has been previously reported within the Dp16 model, as well as in other mouse models of DS^31,45^. Notably though, ALB levels were depleted in the plasma of Dp16 animals relative to WT animals (**Figure 2E, Table S2E**). Although they did not reach significance, SAA showed a trends towards increased levels, while IGF1 and CHI3L1 also showed a trend towards decreased levels in the plasma of Dp16 animals (**Figure S2D, Table S2F,G**).

We have shown previously that the Dp16 mouse model exhibits a liver phenotype characterized by an increase in liver histopathological scoring compared to WT littermates.^46^ We reproduced these results in a new cohort of adult mice, showing that adult Dp16 mice have an increase in overall histopathology scoring of the liver, with the most drastic score coming from the reactive changes parameter (**Figures 2F and 2G, Table S2H)**. This score was almost entirely driven by a so-called ‘ductular reaction’ characterized by proliferation of bile ducts (also known as biliary hyperplasia), increased thickness and number of arteries, and hyperplasia of portal triad endothelial cells (**Figures 2F and G**). Adult Dp16 animals also had elevated levels of inflammation in the liver, with periportal inflammation and accumulation of lymphocytes as contributing features (**Figures 2F and 2G**). To understand the developmental trajectory of this phenotype, we also analyzed liver pathology in a cohort of one-month-old animals (**Figure S2E,F**). Indeed, liver pathology was present as early as one month of age, with modest contributions to the overall pathology score coming from all three categories: cell injury, inflammation, and reactive changes (**Figure S2F, Table S2I**). However, only the elevated reactive changes scores were significant in Dp16 livers at one month of age, suggesting that reactive changes could precede inflammation in the liver. Notably, we did not observe steatosis in either one month or four-month-old animals (**Figures S2F and S2G**). Finally, analysis of canonical plasma markers of liver function in one-month-old animals showed a trend towards elevated alkaline phosphatase (ALP) and creatinine (CREA) in Dp16 animals and decreased SAA and CHI3L1 concentration, while no difference in CRP nor ALB was observed between the two genotypes (**Figure S2H**, **Table S2**K-P).

### Dp16 livers exhibit stellate cell activation and fibrosis

Motivated by our observations of liver pathology in the Dp16 model, we performed transcriptome analysis of the adult Dp16 liver using bulk RNA-sequencing. This analysis revealed around 1500 differentially expressed genes (DEGs) in the Dp16 liver compared to WT (**Figure 3A, Table S3A**). Next, to further explore which pathways may be contributing to liver dysfunction in this model, we performed GSEA, which revealed positive enrichment scores for the Epithelial-Mesenchymal Transition gene set, Inflammatory Response, Interferon Gamma Response, and Coagulation, among others (**Figures 3B, Table S3B**). Negatively enriched pathways included Cholesterol Homeostasis, Bile Acid Metabolism, Fatty Acid Metabolism, and Adipogenesis, all central to liver function (**Figures 3A and 3B**). Within the Epithelial-Mesenchymal Transition pathway, we found significant upregulation of several collagen genes, as well as several genes involved in extracellular matrix remodeling (**Figure 3C**). This includes *Col1a1,* a gene that encodes for alpha chains of type I collagen (**Figure 3D**). Collagen is deposited in excess during injury or disease, causing fibrosis and tissue stiffening.^47^

**Figure 3.**
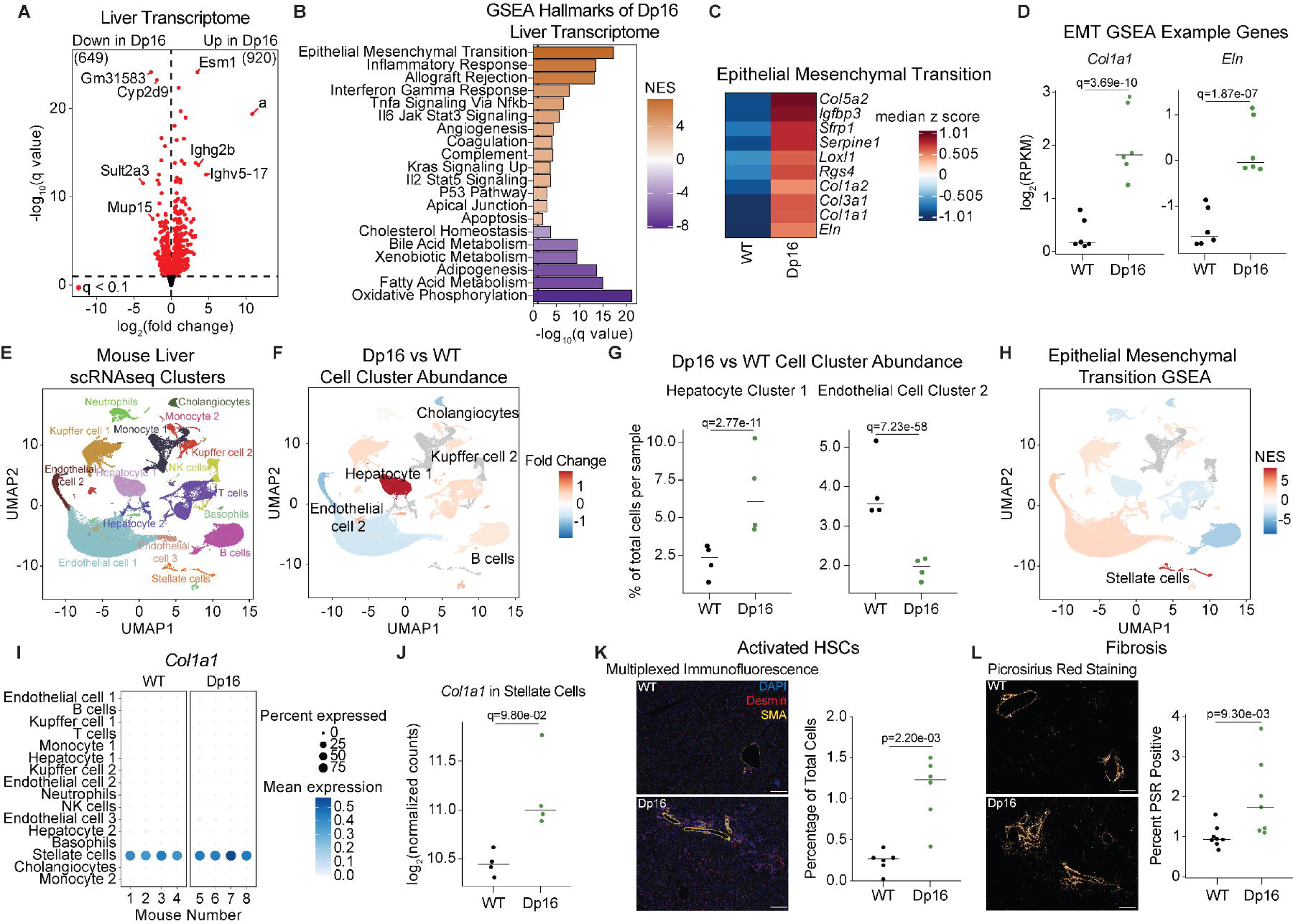
Stellate cell dysfunction drives liver fibrosis in the Dp16 model. (A) Volcano plot summarizing results of whole liver transcriptome analysis of WT (n=6, 3 females) versus Dp16 (n=6, 3 females) mice. Significantly differentially expressed (q<0.1) genes are colored in red. (B) Barplot summarizing the results of Gene Set Enrichment Analysis (GSEA) of gene expression changes in the liver of Dp16 mice (n=6, 3 females) versus wildtype (n=6, 3 females). (C) Heatmap showing the median z-score of the top 10 leading-edge genes from the Epithelial-Mesenchymal Transition gene set. Color coding indicates a positive (red) or negative (blue) median z-score. (D) Sina plots displaying the normalized relative expression (RPKM) of *Col1a1* and *Eln* in both WT and Dp16 mice. Benjamini-Hochberg adjusted *p-value*s (q-values) are indicated. Individual data are presented with a bar at the median. (E) Uniform Manifold Approximation and Projection (UMAP) plot of single cell RNA sequencing analysis of mouse liver, color coded by 16 major cell clusters, identified using Seurat. (F) UMAP plot displaying differential cellular abundance (beta regression Dp16 versus WT) of clusters from single cell RNA sequencing analysis of mouse liver. Significant (FDR < 0.1) clusters are colored according to mean fold-change. (G) Sina plots showing the Dp16 versus WT abundance of Hepatocyte Cluster 1 and Endothelial Cell Cluster 2 from single cell RNA sequencing analysis of mouse liver. Benjamini-Hochberg adjusted *p-value*s (q-values) are indicated. Individual data are presented with a bar at the median. (H) UMAP plot displaying the Epithelial-Mesenchymal Transition gene set Normalized Enrichment Score (NES) for each cluster. (I) Bubble plot summarizing *Col1a1* single-cell expression across clusters and animals, with color representing mean expression and size representing percent of cells with expression. (J) Sina plot displaying the normalized pseudobulk counts of *Col1a1* expression within the Stellate Cell cluster in both WT and Dp16 animals. Benjamini-Hochberg adjusted *p-value*s (q-values) are indicated. (K) Representative images of multiplexed stained liver sections from wildtype and Dp16 mice where DAPI is stained in blue, desmin in red, and smooth muscle actin (SMA) in yellow. Sina plot shows number of desmin+, SMA+ cells as a percent of total cells for wildtype mice (n=6, 3 females) and Dp16 mice (n=6, 3 females). Individual data are presented with a bar at the median and the p-value, as determined by a Mann-Whitney U test, is shown. Scale bars are 100 microns. (L) Representative images of picrosirius red stained (PSR) liver sections from wildtype (n=8, 4 females) and Dp16 (n=7, 4 females) mice, imaged under polarized light. Sina plot showing the percent positive PSR in wildtype and Dp16 mice. Individual data are presented with a bar at the median and the p-value, as determined by a Mann-Whitney U test, is shown. Scale bars are 100 microns.

To begin understanding the contributions of the various cell types in the mouse liver to the observed liver dysfunction, we performed scRNAseq of the WT and Dp16 liver. Clustering and annotation of this data revealed 16 distinct cell clusters representing all major cell types of the liver (**Figures 3E and S3A, Table S4A-R**). When comparing the two genotypes, differential cellular abundance was only observed for a few clusters (**Figure 3F**), with an increase in abundance in the Dp16 liver of hepatocyte cluster 1, as well as increases in Kupffer cell cluster 2 and B cells. Conversely, we observed slight decreases in abundance in endothelial cell cluster 2 and cholangiocytes (**Figures 3F and 3G**). Next, we performed DESeq analysis of cluster by gene-level pseudobulk count data to identify differentially expressed genes within each cluster, followed by GSEA. We observed widespread effects of Dp16 on gene expression across clusters with some gene sets consistently activated (e.g., Allograft Rejection), some activated in a cell type-specific fashion (e.g., Hedgehog Signaling), and others discordant across cell types (e.g., Oxidative Phosphorylation) (**Figure S3B**). As the Epithelial-Mesenchymal Transition gene set was dysregulated in the Dp16 liver bulk transcriptome, we next asked how this pathway was perturbed in specific cell types by plotting the NES values on the UMAP plot. We observed dysregulation of the Epithelial-Mesenchymal Transition pathway in numerous cell types, being activated in some and inactivated in others (**Figure 3H**). Importantly, hepatic stellate cells (HSCs) displayed the highest degree of Epithelial-Mesenchymal Transition activation (**Figure 3H**). When looking at the gene expression of the top-ranked genes for this pathway, for example *Col1a1*, we observed that expression and upregulation was restricted to HSCs (**Figure 3I and 3J**).

To define the contribution of HSCs to the observed Dp16 liver pathology, we next performed multiplexed immunofluorescence analysis using a panel of liver cell type markers. This panel allowed us to visualize major non-immune cell types in the liver, including hepatocytes, cholangiocytes, liver endothelial cells (LECs), liver sinusoidal endothelial cells (LSECs), as well as both inactive and activated HSCs. Consistent with our scRNAseq data, we observed significant increases in abundance of several cell types, one being activated HSCs, specifically around the portal vein (**Figures 3K and S3E, Table S4S**). One key feature of activated HSCs is their ability to produce collagen and other pro-fibrotic factors. To test for collagen buildup in the liver, we stained liver sections with picrosirius red and observed a significant increase in the amount of collagen 1 and 3 in the adult Dp16 liver compared to WT (**Figure 3L**). However, an increase in collagen 1 and 3 was not seen in the one-month-old animals (**Figure S3F**). We also saw a trend towards increased abundance of non-activated HSCs in the Dp16 liver, and a significant increase in both Lyve1+ (lymphatic vessel endothelial hyaluronan receptor-1) and PDPN+ (Podoplanin) cells (**Figures S3B-S3E**). Lyve1 marks LSECs and LECs in the liver and PDPN marks cholangiocytes and lymphatic endothelial cells. Taken together, these data suggest that the activated HSCs are driving the observed Epithelial-Mesenchymal Transition signature in the whole Dp16 liver analysis, and are likely contributors to the observed liver fibrosis, as well as the ductular reaction, in the Dp16 animals.

### Increased copy number of the interferon receptor locus is not a major contributor to Dp16 liver dysfunction

We and others have shown that T21 consistently activates the interferon (IFN) response, driven at least in part by triplication of four of the six interferon receptors (IFNRs) that are encoded on chr21.^4,33^ The Dp16 mouse model harbors a triplication of the four MMU16 encoded *Ifnrs* and it has been shown that dysregulated IFN signaling in this model contributes to various pathophysiological features, including heart defects, developmental delays, cognitive deficits, craniofacial abnormalities and immune hypersensitivity.^48^ Indeed, we observed GSEA signatures of Interferon Gamma Response, along with activation of additional inflammatory signatures, in the Dp16 liver bulk transcriptome analysis (**Figures 4A and S4A**). Dp16 livers strongly upregulate some genes in the Interferon Gamma Response, including *Cxcl9*, which encodes an IFN-stimulated cytokine that attracts immune cells to sites of infection (**Figure 4B**). Given the important role of increased IFN signaling in various DS phenotypes, we next asked which cell types in the liver exhibit dysregulation of this pathway at the single-cell level. Interferon Gamma Response dysregulation was seen in almost every cell type, with the exceptions of endothelial cell cluster 2 and cholangiocytes (**Figure 4C**). The greatest upregulation was seen in neutrophils, monocytes, and in the two Kupffer populations (**Figure 4C**). *Cxcl9* expression was seen to be most significantly dysregulated in endothelial cell cluster 1, with values in Dp16 mice that completely discriminated the group from WT animals (**Figures 4D and 4E**).

**Figure 4.**
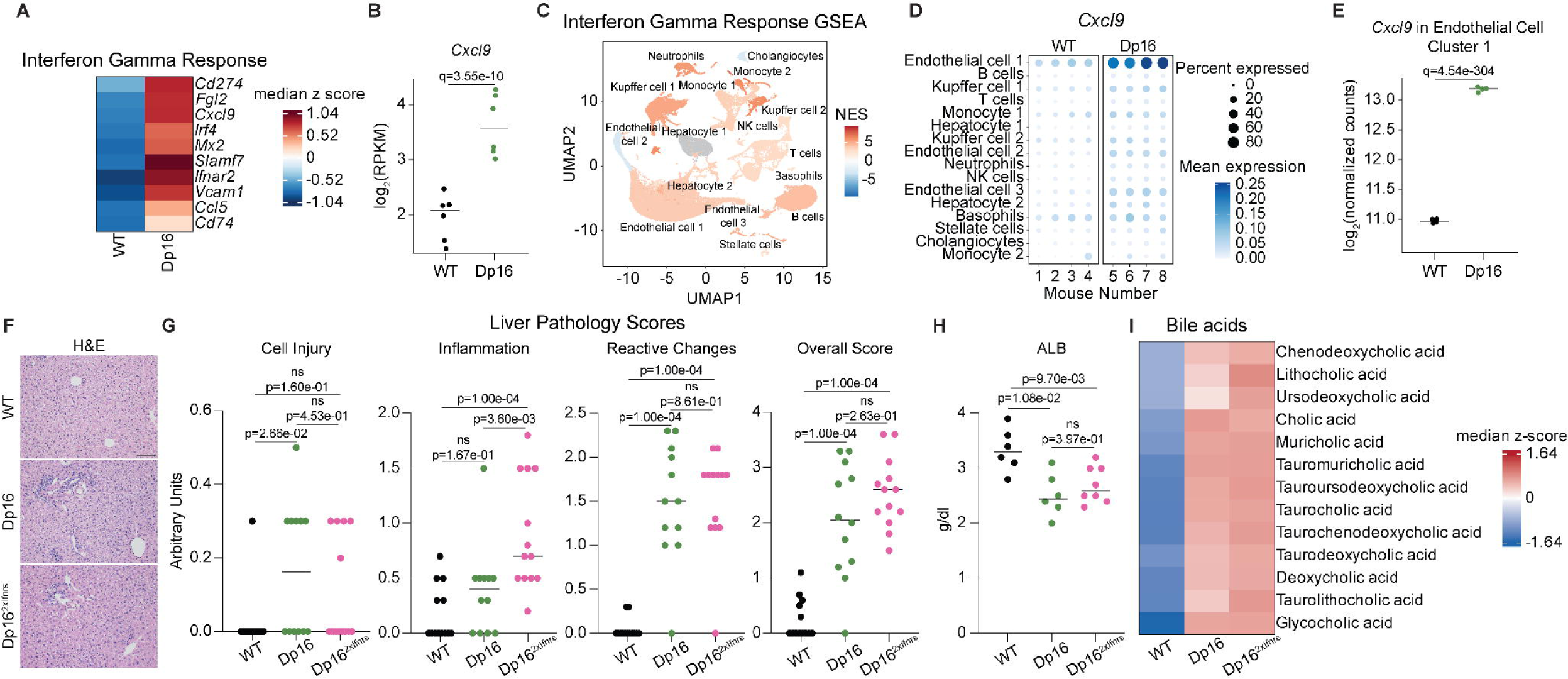
Increased copy number of the interferon receptor locus is not a major contributor to Dp16 liver dysfunction. (A) Heatmap showing the median z-score of the leading-edge genes from the Interferon Gamma Response gene set from WT and Dp16 liver bulk transcriptome analysis. (B) Sina plot displaying relative expression (RPKM) of *Cxcl9* in both wildtype and Dp16 liver bulk transcriptome analysis. Benjamini-Hochberg adjusted *p-value*s (q-values) are indicated. (C) UMAP plot displaying the Interferon Gamma Response gene set normalized enrichment score (NES) for each cluster. (D) Bubble plot displaying the mean expression and the percent of cells per cluster expressing *Cxcl9* in both wildtype and Dp16 mice. (E) Sina plot displaying the log_2_ of the normalized counts of *Cxcl9* expression within Endothelial Cell cluster 1 in both wildtype and Dp16 animals. Benjamini-Hochberg adjusted *p-value*s (q-values) are indicated. Individual data are presented with a bar at the median. (F) Representative images of H&E-stained liver sections from adult wildtype, Dp16, and Dp16^2xIFNRs^ mice. Scale bar is 100 microns. (G) Sina plots showing the metrics of liver pathology scoring from adult wildtype (n=13, 6 females), Dp16 (n=12, 5 females), and Dp16^2xIFNRs^ (n=13, 7 females) mice. Individual data are presented with a bar at the median and p-values, as determined by a Mann-Whitney U test, are shown. (H) Sina plot displaying the grams/deciliter of plasma Albumin from adult wildtype (n=6, 4 females), Dp16 (n=6, 2 females), and Dp16^2xIFNRs^ (n=8, 1 female) mice. Individual data are presented with a bar at the median and p-values, as determined by a Mann-Whitney U test, are shown. (I) Heatmap displaying the median z-score of various plasma bile acids in adult wildtype (n=9, 4 females), Dp16 (n=13, 6 females), and Dp16^2xIFNRs^ (n=10, 5 females) mice.

To examine the contribution of increased gene dosage of the *Ifnrs* in the liver pathology of DS, we employed the Dp16^2xIfnrs^ mouse model.^48^ This modified Dp16 mouse model has normalized copy number of the four *Ifnrs;* from three copies back to two, while maintaining the rest of the segmental duplication, allowing us to define the contribution of the triplication of these receptors. Histopathological analysis of these mice revealed that normalization of the *Ifnrs* did not rescue the liver pathology observed in adult Dp16 mice (**Figures 4F and 4G, Table S5A**). At four months of age, the Dp16^2xIfnrs^ animals surprisingly had significantly higher inflammation scores compared to both WT and Dp16 mice and trended towards increased reactive changes and overall scores (**Figure 4G**). At one month of age, Dp16^2xIfnrs^ scores trend lower than Dp16 scores, but none reached significance (**Figures S4B and S4C, Table S5B**). These observations may be due to the proposed anti-inflammatory properties of one of the normalized receptors, IL10RB, and its respective cytokine, IL10.^49,50^ We next tested the impact of the *Ifnr* copy number normalization on other features observed in the adult Dp16 model. Dp16^2xIfnrs^ animals did not have increased plasma albumin abundance compared to Dp16 animals (**Figure 4H, Table S5C**), nor did they have decreased liver fibrosis compared to Dp16 animals (**Figure S4D**). One month old Dp16^2xIfnrs^ also did not have decreased levels of fibrosis compared to Dp16 animals (**Figure S4E, Table S5D)**. Importantly though, Dp16^2xIfnrs^ animals retained elevated levels of plasma BAs (**Figure 4I, Table S5E-F**). Overall, these data suggest that the increased *Ifnr* gene dosage in the Dp16 mouse model is not a major driver of the observed liver pathology and BA metabolism dysregulation.

### Metabolic dysfunction in the Dp16 liver is associated with specific cell populations

Next, we turned our attention towards an interrelated group of metabolic signatures identified within the bulk liver transcriptome as being negatively enriched in Dp16 compared to WT, including Cholesterol Homeostasis, Adipogenesis, Bile Acid Metabolism, and Fatty Acid Metabolism (**Figures 3A and 5A**). Within the Cholesterol Homeostasis Pathway, significantly downregulated genes include Farnesyl Pyrophosphate (*Fdps*) and NAD(P) Dependent Steroid Dehydrogenase-like (*Nsdhl*) (**Figure 5B)**. *Fdps* is involved in the later stages of the cholesterol biosynthesis pathway, producing a key cholesterol precursor, farnesyl pyrophosphate.^47^ *Nsdhl* encodes an enzyme responsible for converting lanosterol into cholesterol and has been shown to be important for the transport of lipid droplets.^48^ Within the Bile Acid Metabolism signature, two significantly downregulated leading-edge genes include Solute Carrier Family 23 member 1 (*Slc23a1*) and Cytochrome P450 46A1 (*Cyp46a1)* (**Figure 5B**). *Slc23a1*, which encodes the sodium-dependent vitamin C transporter protein, is important for maintaining vitamin C levels in the body.^52^ These data are consistent with a strong impact of Dp16 gene triplication on hepatic metabolism.

**Figure 5.**
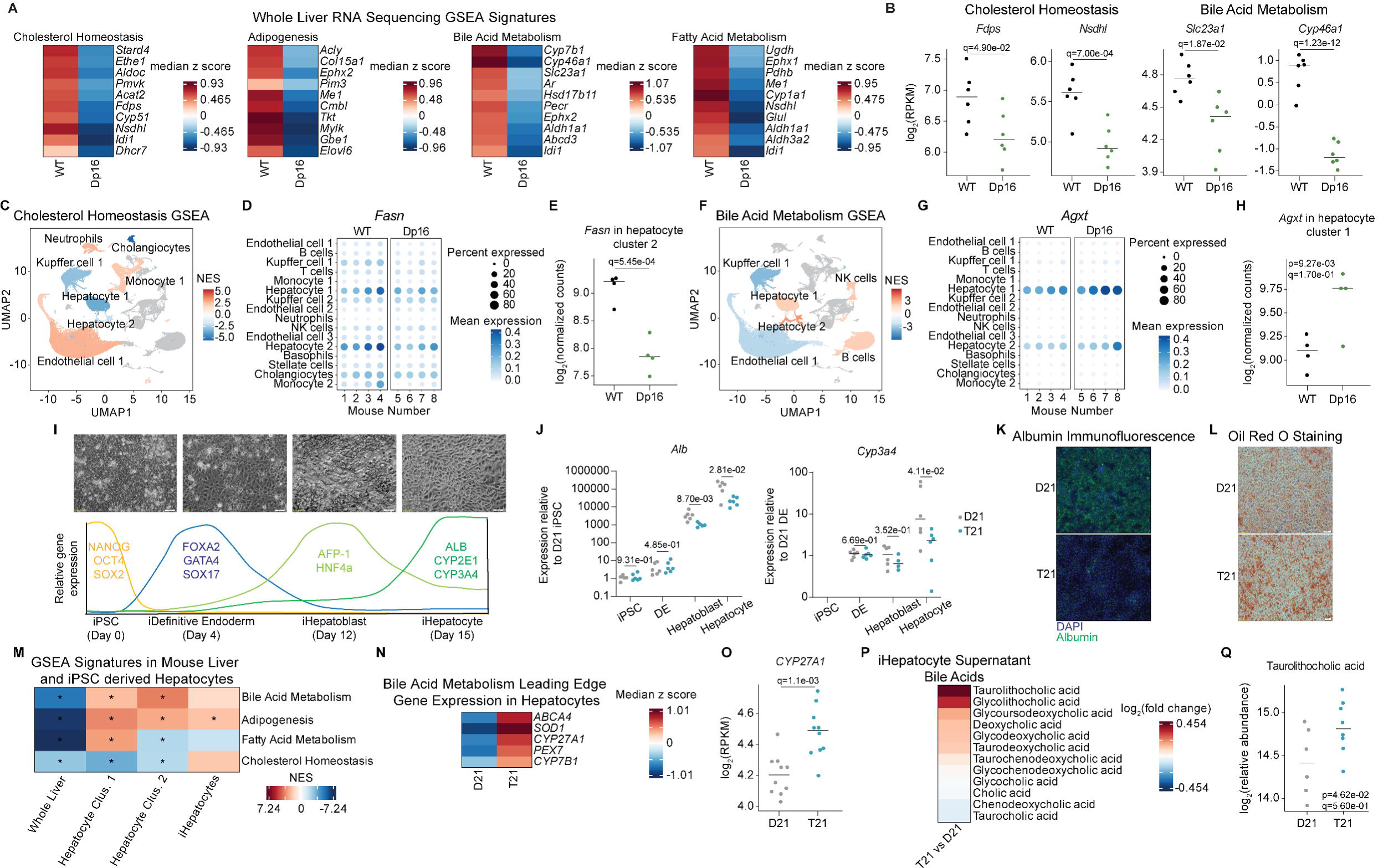
Liver dysfunction in Down syndrome is driven by hepatocyte dysfunction. (A) Heatmaps showing the median z-score of the leading-edge genes from the Cholesterol Homeostasis, Adipogenesis, Bile Acid Metabolism, and Fatty Acid Metabolism GSEA signatures in whole liver tissue. Color coding indicates a positive (red) or negative (blue) median z-score. (B) Sina plots displaying the log_2_(RPKM) of various genes. Benjamini-Hochberg adjusted *p-value*s (q-values) are indicated. (C) UMAP plot displaying the Cholesterol Homeostasis GSEA signature enrichment in all clusters. Color coding indicates a positive (red) or negative (blue) normalized enrichment score. (D) Bubble plot displaying the mean expression and the percent of cells per cluster expressing *Fasn* in both wildtype and Dp16 mice. (E) Sina plot displaying the log_2_ of the normalized counts of *Fasn* expression within hepatocyte cluster 2 in both wildtype and Dp16 animals. The q-value indicates linear regression significance after Benjamini-Hochberg correction for multiple testing. (F) UMAP plot displaying the Bile Acid Metabolism gene set normalized enrichment score (NES) in each cluster. Color coding indicates a positive (red) or negative (blue) NES. (G) Bubble plot displaying the mean expression and the percent of cells per cluster expressing *Agxt* in both wildtype and Dp16 mice. (H) Sina plot displaying the log_2_ of the normalized counts of *Agxt* expression within hepatocyte cluster 2 in both wildtype and Dp16 animals. The q-value indicates linear regression significance after Benjamini-Hochberg correction for multiple testing. (I) Schematic illustrating the genes measured at various stages in the iPSC derived hepatocyte differentiation protocol, as well as microscopy images of cells in each stage. Scale bars are 50 microns. (J) Sina plots displaying the gene expression of *Alb* and *Cyp3a4* relative to disomic iPSCs. Values from disomic cells are shown in gray and from trisomic cells are shown in blue. Individual data are presented with a bar at the median and p-values, as determined by a Mann-Whitney U test, are shown. (K) Representative images of Albumin (green) and DAPI (blue) immunofluorescent staining of disomic and trisomic iPSC derived hepatocytes. (L) Representative images of Oil Red O staining of disomic and trisomic iPSC derived hepatocytes. Scale bars are 50 microns. (M) Heatmap displaying the enrichment of various GSEA gene sets in whole mouse liver, hepatocytes, and two single cell hepatocyte clusters. Asterisks indicate significance after Benjamini-Hochberg correction for multiple testing (q-value< 0.1). (N) Heatmaps showing the median z-score of the leading-edge genes from the Bile Acid Metabolism GSEA pathway in the iHepatocytes. (O) Sina plots displaying the log_2_(RPKM) of *CYP27A1*. Benjamini-Hochberg adjusted *p-value*s (q-values) are indicated. (P) Heatmap displaying the log_2_(fold change) of bile acids detected in the supernatant of trisomic versus disomic iPSC derived hepatocytes. Color coding indicates a positive (red) or negative (blue) log_2_(fold change). (Q) Sina plot displaying the log_2_(relative abundance) of Taurolithocholic acid detected in the supernatant of disomic and trisomic iPSC derived hepatocytes. Benjamini-Hochberg adjusted *p-value*s (q-values) are indicated. P-value was determined by a Mann-Whitney U test.

We then aimed to understand which cell types are associated with this observation of metabolic dysfunction in the whole tissue by analyzing these signatures within our scRNAseq data. We first analyzed the Cholesterol Homeostasis signature and observed it is negatively enriched in several clusters, including both hepatocyte clusters, cholangiocytes, and Kupffer cells (**Figure 3C**). We observed that *Fasn*, which encodes fatty acid synthase, is most significantly dysregulated in the same clusters which exhibit the greatest perturbations in the Cholesterol Homeostasis signature: cholangiocytes, the hepatocyte clusters, and Kupffer cells (**Figures 5D and 5E**). These data suggest dysregulated cholesterol homeostasis in the whole Dp16 liver is driven by a few specific cell types, including hepatocytes.

We next analyzed the Bile Acid Metabolism signature within our single cell data and observed that this signature is highly discordant across cell types, despite being strongly downregulated in the bulk transcriptome data (**Figures 5A and 5F**). Whereas this signature is negatively enriched in endothelial cells, for example, it is positively enriched in both hepatocyte clusters (**Figures S3B and 5F**). Hepatocyte specific perturbations are apparent in the top-ranked genes, including *Agxt* and *Apoa1* (**Figures 5G, 5H, S5A, S5B**). *Agxt* encodes for alanine-glyoxylate aminotransferase. This enzyme plays a role in glyoxylate detoxification in the peroxisome, the main site of BA metabolism in the cell, thereby maintaining the overall health of this important organelle. *Apoa1* encodes Apolipoprotein A1, which is a major component of high density lipoproteins (HDL) and, paradoxically, can be downregulated by BAs via FXR.^51^ These data suggests that the Dp16 model exhibits intrinsic deficits in maintaining cholesterol, fatty acid, and BA homeostasis and that these deficits are driven primarily by dysfunctional hepatocytes.

Given the impact of hepatocyte function on the overall health of the liver, we next aimed to study the cell-intrinsic features of hepatocyte dysfunction. Towards this end, induced pluripotent stem cells (iPSCs) were differentiated into hepatocyte-like cells (iHepatocytes) using a panel of iPSCs from individuals with T21 and controls. Briefly, iPSCs were differentiated into the definitive endoderm (iDE) stage, then the iHepatoblast stage, which were then matured into iHepatocytes using a combination of dimethyl sulfoxide (DMSO), hepatocyte growth factor (HGF), and dexamethasone (dex) (**Figure S6A**).^52^ Successful differentiation was tracked using quantitative real-time PCR (qRT-PCR) to measure the expression of stage-specific markers (**Figure 5I, Table S6A,B**). Gene expression measurements revealed reduced expression of mature hepatocyte genes *ALB* and *CYP3A4* in T21 iHepatocytes when compared to euploid controls (**Figure 5J)**. Diminished albumin levels in T21 iHepatocytes were confirmed by immunofluorescence staining (**Figure 5K).** Oil Red O staining for lipid accumulation, another feature of hepatocyte function, revealed elevated lipid storage in T21 iHepatocytes compared to controls (**Figure 5L**).

Transcriptome analysis of T21 iHepatocytes revealed hundreds of DEGs both up- and downregulated relative to euploid controls (**Figure S5B, Table S6C**). GSEA of this transcriptome data was dominated by down-regulation of cell cycle-related gene sets such as E2F Targets, G2M Checkpoint and Myc Targets V1, which could be reflective of diminished proliferative capacity in the T21 iHepatocytes (**Figure S5C**). Next, we compared normalized enrichment scores for the metabolic ‘module’ from the bulk liver transcriptome and single-cell hepatocyte clusters 1 and 2 to the iHepatocytes and observed widespread discordance between the bulk and hepatocyte-specific analyses. For example, BA metabolism and adipogenesis were downregulated in the bulk and upregulated in all three hepatocyte-specific analyses, whereas cholesterol metabolism was downregulated in all but the iHepatocytes (**Figure 5M**). Closer examination of top-ranked genes for BA metabolism in the iHepatocytes revealed strong upregulation of a number of genes shown to be central to this process. This includes *CYP27A1*, which encodes for the enzyme that controls the rate limiting step in BA biosynthesis (**Figures 5N and 5O**).

Finally, given the observed elevation of plasma BAs in individuals with DS and that hepatocytes are the cells which produce these BAs *in vivo*, we next measured BA abundance in both iHepatocytes and supernatants. We observed a trend towards an increase in numerous BAs in the iHepatocytes, as well as in the supernatant (**Figures 5P, 5Q, and S5E-S5G**). Together, these data suggest that T21 is associated with cell-intrinsic dysregulation of hepatocyte function, where expression of key mature hepatocyte genes appears to be diminished, while other hepatocyte features, such as lipid storage, are elevated. The fact that not all features of mature hepatocytes are diminished in T21 iHepatocytes suggests a more complex dysregulation than a simple failure to mature into hepatocyte-like cells.

### Liver pathology in Dp16 mice is dependent on diet

Given that metabolic homeostasis is sensitive to diet, we next investigated the impact of diet on BA metabolism and liver function in Dp16 mice. Animals were fed either a high-fat diet (HFD, 60 kcal% fat) or a control low-fat diet (LFD, 10 kcal% fat with all other components matched to the HFD) for two months beginning at one month of age (**Figure S6A**). As expected, both WT and Dp16 animals on the HFD gained significantly more body weight over the course of treatment than the animals on the LFD, with a trend towards increased liver mass (**Figures S6B and S6C**). Dp16 animals were slightly heavier than their WT counterparts after two months on the LFD, but not the HFD (**Figure S6B**). Notably, Dp16 animals do not develop steatosis or obesity on their standard diet, which contains 16 kcal% fat (Figure S2G). Histopathological analysis of the Dp16 liver revealed a hypersensitivity to diet-induced liver injury, with HFD-fed Dp16 animals exhibiting significant elevations in inflammation and reactive changes, as was the case for animals fed standard chow (**Figures 6A and 6B**). The HFD-fed Dp16 animals, however, also showed a significant elevation in steatosis (**Figure 6B**). Most strikingly, Dp16 animals on the LFD exhibited no elevation in liver pathology relative to their WT counterparts, indicating that liver pathology, even under normal chow conditions, is the product of diet-induced injury (**Figures 6B and S6D**). In addition, none of the treatment groups had a significant increase in fibrosis (**Figure S6E**). Finally, when we broke out our treatment groups by sex, we observed elevated pathology for both sexes at the level of overall score (**Figure S6F**).

**Figure 6.**
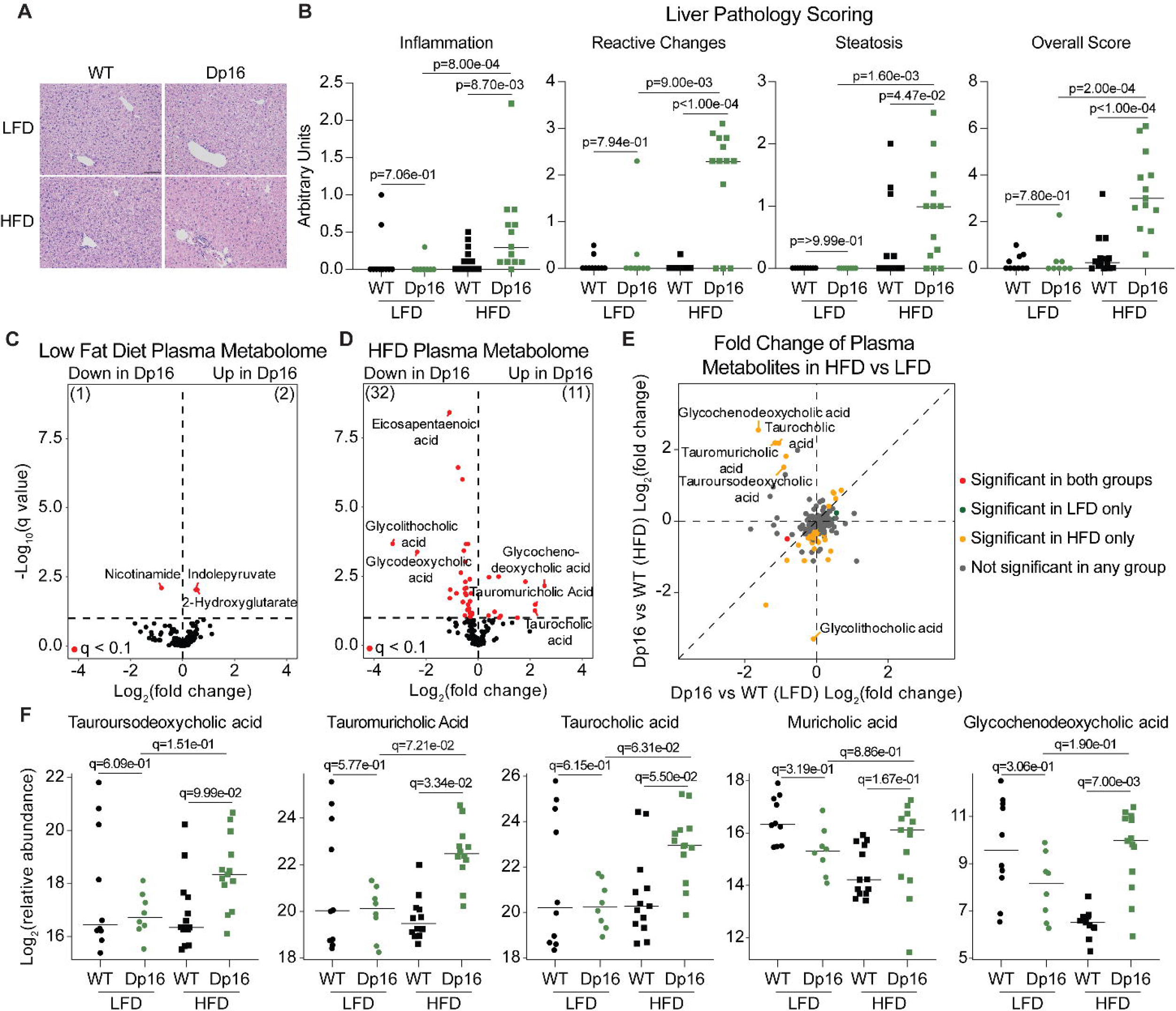
Liver pathology in Dp16 mice is dependent on diet. (A) Representative images of H&E-stained liver sections from wildtype and Dp16 mice fed either a LFD or HFD. Scale bar is 100 microns. (B) Sina plots showing the metrics of liver pathology scoring from WT and Dp16 mice fed either a LFD or HFD (WT LFD n=10, 4 females, Dp16 LFD n=8, 4 females, WT HFD n=14, 4 females, Dp16 HFD n=13, 4 females). Individual data are presented with a bar at the median and p-values, as determined by a Mann-Whitney U test, are shown. (C) Volcano plot summarizing results of plasma metabolomic analysis of Dp16 LFD mice compared to wildtype LFD mice (WT LFD n=10, 4 females, Dp16 LFD n=8, 4 females). Significantly differentially expressed (q<0.1) genes are colored in red. (D) Volcano plot summarizing results of plasma metabolomic analysis of Dp16 HFD mice compared to wildtype HFD mice (WT HFD n=13, 4 females, Dp16 HFD n=13, 4 females). Significantly differentially expressed (q<0.1) genes are colored in red. Benjamini-Hochberg adjusted *p-value*s (q-values) are indicated. (E) Scatterplot comparing fold-changes of plasma metabolites from Dp16 LFD mice compared to WT LFD mice (x axis) versus Dp16 HFD mice compared to WT HFD mice (y axis). Points are colored according to significance in both groups (red), significance in the LFD group only (green), significance in the HFD group only (yellow), or not significant in any group (gray). Significance is defined as q < 0.1 (10% FDR). (F) Sina plots displaying the relative abundance of indicated plasma bile acids. Benjamini-Hochberg adjusted *p-value*s (q-values) are indicated.

Next, we analyzed the plasma metabolome of this cohort and made several interesting observations between different comparison groups. First, we observed that the global dysregulation of the plasma metabolome of Dp16 animals fed the standard chow were almost completely abrogated in the LFD group (**Figures 6C, 6D, and 1F**). Furthermore, HFD-fed Dp16 animals revealed a strong dysregulation in BA metabolism not seen in HFD-fed WT animals (**Figure 6D**). Indeed, direct comparison of dietary impact on the plasma metabolome highlighted this profound impact on BA metabolism in Dp16 animals (**Figures 6E, S6F, S6G, and S6H**). Closer inspection of individual BAs revealed various modes of dysregulation. For example, tauroursodeoxycholic acid, tauromuricholic acid, and taurocholic acid, all BAs that have been conjugated to taurine to facilitate biliary secretion, showed similar patterns: not differentially abundant in LFD-fed animals and elevated only in HFD-fed Dp16 animals (**Figure 6F**). Muricholic acid, in contrast, showed a trend towards lower levels in LFD-fed Dp16 animals compared to LFD-fed WT animals, and was decreased in HFD-fed WT animals, but not HFD-fed Dp16s (**Figure 6F**). Glycochenodeoxycholic acid showed similar trends to muricholic acid, but with a significant difference in the overall impact of HFD between Dp16 and WT animals (**Figure 6F**). Taken together, these data indicate that alterations in BA metabolism and liver pathology are mediated by diet in the Dp16 mouse model of DS.

### Diet shapes the Dp16 hepatic transcriptome

We next analyzed the liver transcriptome of the WT and Dp16 animals under LFD and HFD conditions. When comparing Dp16 to WT under either LFD or HFD conditions, we detected thousands of DEGs (**Figures 7A and S7A**). Comparison of Dp16 chr16-triplicated DEGs demonstrated a strong concordance between the two treatment conditions with 71 of 82 genes (86.6%) present in both analyses (**Figure 7B**). This was not the case, however, when comparing DEGs in the non-triplicated region, where only 748 of 3,531 DEGs (21.2%) were present in both analyses (**Figure 7B**). GSEA of transcriptome data from the various experimental groups demonstrated genome-wide impacts of diet on the transcriptome in both WT and Dp16 mice (**Figures S7B and S7C**). Characteristic signatures including elevated Interferon Signaling and decreased Fatty Acid Metabolism were present regardless of diet (**Figure S7C**). We next compared normalized enrichment scores (NES) from the Dp16 versus WT LFD analysis and the Dp16 versus WT HFD analysis to visualize transcriptomic differences in response to diet. Here, we observed that many gene sets grouped along the x=y axis, suggesting they were perturbed similarly in both diet groups. Numerous gene sets deviated from this pattern, however. Interferon and inflammation related pathways, for example, were activated in LFD conditions and repressed or less activated in HFD conditions, clustering in the bottom right quadrant, (**Figure 7C**). Examination of key top-ranked genes from the Interferon Alpha Pathway demonstrated that this differential response is due to IFN activation in WT mice that is blunted in Dp16 (Dp16 versus WT LFD analysis). This observation could explain the lack of rescue by *Ifnr* copy number normalization (**Figure 4**).

**Figure 7.**
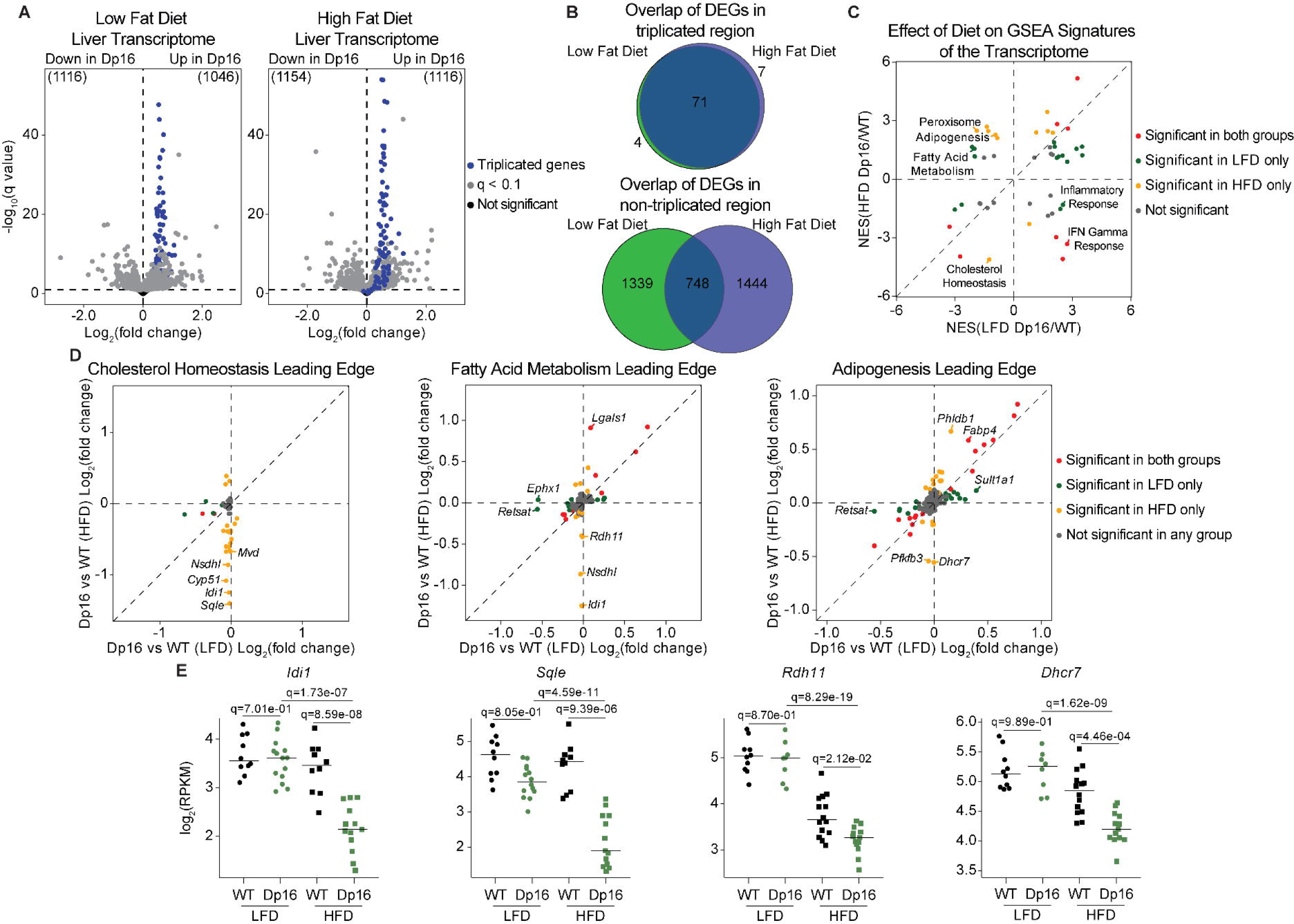
A high-fat diet contributes to global transcriptomic changes in the Dp16 liver. (A) Volcano plots summarizing the results of transcriptome analysis of livers from Dp16 mice fed a low-fat diet (LFD) compared to wildtype mice fed a LFD and from Dp16 mice fed a high-fat diet (HFD) compared to wildtype mice fed a HFD. (wildtype LFD n=10, 4 females, Dp16 LFD n=8, 4 females, wildtype HFD n=14, 4 females, Dp16 HFD n=13, 4 females). Points in blue are in the MMU16 region triplicated in Dp16 mice, points in gray have reached a linear regression significance threshold of less than 0.1 after Benjamini-Hochberg correction for multiple testing, and points in black are not significant. (B) Venn diagrams displaying the overlap in the number of significant differentially expressed genes in the Dp16 triplicated region between HFD fed mice and LFD mice (top) or in the differentially expressed genes that are not triplicated in Dp16 mice (bottom). (C) Scatter plot comparing GSEA NES values for signatures in the plasma of Dp16 versus WT mice on LFD (x axis) and Dp16 versus WT mice on HFD diet (y axis). Points are colored according to significant in both groups (red), significant in the LFD group only (green), significant in the HFD group only (yellow), or not significant in either comparison (gray). Significance is defined as *q* < 0.1. (D) Scatter plots comparing the fold-changes of leading-edge genes from the cholesterol homeostasis, fatty acid metabolism, and adipogenesis Hallmark gene sets for Dp16 versus WT mice on LFD (x axis) and Dp16 versus WT on HFD (y axis). Points are colored according to significant in both groups (red), significant in the LFD group only (green), significant in the HFD group only (yellow), or not significant in any group (gray). Significance is defined as *q* < 0.1. (E) Sina plots displaying the log_2_(RPKM) of various genes. Horizontal bars indicate median values. Benjamini-Hochberg adjusted *p-value*s (q-values) are indicated.

Other prominent gene sets that showed diet-specific effects include Peroxisome, Adipogenesis, Fatty Acid Metabolism, and Cholesterol Homeostasis. The Peroxisome and Adipogenesis gene sets were repressed under LFD conditions and significantly activated under HFD conditions. Cholesterol Homeostasis was modestly, but not significantly, repressed by LFD but strongly repressed by HFD (**Figure 7C**). Next, we compared the effect of diet and genotype on the behavior of various top-ranked genes from metabolism-related gene sets of interest. We found that very few of the top-ranked genes within the Cholesterol Homeostasis gene set were differentially expressed in both HFD and LFD Dp16 animals relative to WTs. Despite the gene sets not rising to the level of significance at the NES level, several leading-edge genes were significantly downregulated in Dp16 animals under LFD conditions, consistent with a defect in cholesterol homeostasis even in the absence of a fatty diet (**Figure 7D**). Conversely however, numerous unique top-ranked genes were downregulated in Dp16 animals upon HFD, indicating a differential response to diet (**Figure 7D**). Among those genes significantly downregulated in HFD-fed Dp16 animals were *Idi1* and *Sqle*, both of which are required for cholesterol biosynthesis, with *Sqle* controlling the rate-limiting step in the process (**Figure 7E**).

Fatty Acid Metabolism genes showed similar patterns to those in Cholesterol Homeostasis, with genes downregulated in LFD-fed Dp16 that varied from those downregulated under HFD conditions (**Figure 7D**). For example, *Retsat* and *Ephx1*, two genes involved in lipid metabolism, were both downregulated only under LFD-fed conditions, which could contribute to an altered response to HFD (**Figure 7D and S7D**). *Nsdhl*, which was downregulated in our four-month-old cohort fed standard chow, was only downregulated in Dp16 by HFD (**Figure 7D**). Adipogenesis related genes were variably impacted by diet. For example, *Fabp4*, which binds to fatty acids and assists with uptake and metabolism was upregulated in Dp16 liver by either diet (**Figure 7D**). Notably, among the genes only affected by HFD in this group was *Dhcr7*, which encodes the enzyme that catalyzes the final step in cholesterol biosynthesis (**Figures 7D**). Finally, BA metabolism was upregulated by HFD regardless of genotype, with a slightly greater elevation in Dp16 animals (**Figure S7B**) Taken together with our standard chow fed Dp16 data, these observations are consistent with a broad dysregulation of metabolism in Dp16 animals that is greatly exacerbated by a HFD and rescued by a LFD.

## DISCUSSION

Here we report that both individuals with T21 and the Dp16 mouse model of DS exhibit elevated levels of plasma BAs at baseline. We show that this is accompanied by plasma proteomic biosignatures consistent with liver dysfunction in the human population with DS. The Dp16 mouse model exhibits aberrant liver histopathology at baseline,^46^ an observation which we replicate here. Using histopathological assessment and multicolor immunofluorescence, we demonstrate that the pathology is typified, not only by inflammatory infiltrates, but by portal triad endothelial hyperplasia, increased numbers of thickened arteries, and ductular reaction. Transcriptome analyses revealed that Dp16 livers are marked by elevated gene expression programs for interferon signaling and inflammation as well as depressed metabolic programs associated with cholesterol and fatty acid metabolism. We show that normalization of the copy number for the four interferon receptors on mouse chr16 does not rescue liver pathology in these mice. Dp16 mice showed a heightened sensitivity to diet-induced liver injury, characterized by increased pathology – including steatosis, elevated plasma BAs, and widespread impacts on the transcriptome. Further studies will be needed to define the extent to which these mechanistic investigations in mice hold true in humans, however, we show that human iPSC-derived hepatocytes from individuals with T21 exhibit cell-intrinsic defects in metabolic gene expression programs and elevated BA levels.

Extensive research in humans and animals has demonstrated widespread consequences for altered liver function including renal toxicity,^53^ neurological impairment,^54^ pulmonary inflammation,^55^ and even reproductive deficits.^56^ While it is known that T21 impacts essentially every organ and tissue system in the body, it is unclear whether or how altered liver function might contributes to or exacerbate these conditions. The potential links between altered BA metabolism, obesity, and metabolic syndromes in T21 are readily apparent though.^17,57–59^ The impact of liver dysfunction on IGF1 levels, of which we have shown to be decreased in T21 individuals, alone could have stark consequences for development.^9^ We recently demonstrated that levels of the liver-derived protein IGF1 in the population with DS are: 1) lower across the lifespan, 2) inversely correlated with neurodegeneration markers, including Neurofilament Light (NfL), 3) inversely correlated with inflammatory markers such as TNFα, and 4) associated with short stature.^9^ These results represent just one way in which altered BA metabolism and hepatic protein synthesis could affect multiple systems within DS. Indeed, liver disease in the typical population has been linked to numerous conditions that are more prevalent in the population with DS, including congenital heart defects,^60^ celiac disease,^61^ hypothyroidism,^62^ alopecia areata,^63,64^ and even Alzheimer’s disease.^65^

Liver disease in the typical population is often accompanied by metabolic abnormalities, such as dyslipidemia, hypoalbuminemia, hypoglycemia, and dysregulated BA metabolism. This, combined with increased body mass index (BMI) levels, can exacerbate certain liver diseases. Dyslipidemia, high levels of lipids in the blood, is more common in the population with DS and even more so in those that are obese.^66^ Although there is some evidence to suggest that obesity is a driver of dyslipidemia in DS^67,68^, other reports show that dyslipidemia occurs regardless of weight status.^66,69^ However, the multitude of potential contributing features (diet, obesity, genetic factors, among others) to metabolic alterations in people with DS makes it hard to disentangle the root cause. Here, we provide evidence that, independent of BMI, individuals with DS show clear elevations plasma BAs, consistent with liver dysfunction as does the Dp16 mouse model of DS and that these changes can be modified by diet. These data support the notion that there are intrinsic differences contributing to metabolic alterations in individuals with DS that are independent of their BMI, and that these alterations are exacerbated by a diet high in fat in a Down syndrome mouse model.

BAs are derived from cholesterol in hepatocytes and play many roles, mainly working as emulsifiers for the absorption of fats and signaling molecules for various pathways. Given their diverse biological roles, there are potentially wide-ranging repercussions for dysregulated BA metabolism and signaling. For instance, associations between altered BA metabolism and the pathogenesis of various disorders, such as cardiometabolic and inflammatory diseases, have been documented.^18^ BAs can bind to receptors expressed on innate immune cells, such as macrophages and dendritic cells, and impact immune cell function, modulate inflammation, potentially contributing to the development of long-term immune-related conditions. ^70^ Increased activation of the IFN pathway has been shown to contribute to many DS phenotypes in both animal models of Down syndrome and iPSC derived cell types.

Dysregulated BA metabolism can also impact lipid metabolism. While baseline expression of BA receptors is crucial for maintaining proper lipid metabolism, increased activation of BA receptors through increased levels of BAs disrupts lipid homeostasis and leads to increased lipid storage in the liver.^18^ A diet high in fat can significantly disrupt the BA pool and increase the secretion of BAs, showing the influence that diet plays on the delicate balance of this pool.^71^ Our data specifically shows a strong connection between a HFD (60 kcal%), increased levels of plasma BAs, and liver steatosis, which is further corroborated by data showing that a diet lower in fat (10 kcal%) can reverse this steatosis and return plasma BAs to a more normal level. Importantly, animals fed their standard diet (16 kcal% fat) do not develop steatosis yet have dysregulated BA metabolism.

We have also shown here that Dp16 mice fed their standard diet exhibit liver pathology characterized by an increase in many features, mainly inflammation and ductular reaction. It is not surprising that we observe liver inflammation, as a plethora of factors can lead to liver inflammation, including BA dysregulation.^72^ Ductular reaction is the proliferation of bile ducts, often in response to injury or inflammation. It is well established that this phenotype arises when hepatic progenitor stem cells become activated upon liver injury, and more recent research has shown that ductular reaction specifically occurs to reconstruct the biliary tree network by creating new bile secreting ducts.^73^ The ductular reaction in our mouse model is likely a response to diet induced injury, as this phenotype is not seen in the animals fed a low-fat diet. The ductular reaction phenotype is also likely tied to dysregulated BA metabolism in our model, as damaged bile ducts have been shown to lead to elevated plasma BAs, and – again – LFD also ameliorates the observed elevated plasma BAs.^22^ Further experiments will need to be performed though to fully understand the cause-and-effect relationship at play.

Bulk transcriptome analysis identified profound impacts of Dp16 mice on global gene expression under all dietary conditions, with effects spanning inflammation, fibrosis (Epithelial-Mesenchymal Transition), and metabolism. scRNAseq revealed that some alterations were widespread, impacting nearly all cell types, whereas others were only perturbed in a subset of cell types. Many programs were differentially impacted by genotype across cell types, upregulated in some and downregulated in others. Metabolism fell into the last category, with interesting patterns observed for two clusters of hepatocytes. Indeed, we were able to use human iPSCs to derive hepatocytes, confirming the impact of genotype on gene expression in a cell autonomous fashion. Broadly, we observed that T21 hepatocytes display many of the same deficits observed in both our human and mouse analyses, showing that these phenotypes are cell intrinsic. For example, in the same way as is observed in both our human and mouse plasma analyses, albumin expression is decreased in the differentiated T21 iHepatocytes. Dysfunctional hepatocytes have implications for the overall metabolic function of the liver, as they play key metabolic roles.^74^

### Limitations of the study

One limitation of this study is the use of mice to understand human metabolism. While mice and humans are similar in many regards, they also vary in many ways.^75^ For example, in regard to metabolism, mice and humans differ in the mitochondrial density of their cells, the rate of reactive oxygen species production, their brown fat deposition, and the relative mass of metabolically active tissues, such as the liver.^76^ Additionally, while the Dp16 mouse model is one of the most used models of DS, it does not contain a freely segregating extra chromosome, present in ∼95% of cases with DS.^77^ The Dp16 model also does not confer full trisomy of the human chromosome 21 mouse orthologs, but rather around 120 of the protein coding genes on mouse chromosome 16. Another limitation of this study is the collection of HFD data at a single time point. Here, we show the impact of a HFD on mice fed the diet for 2 months, beginning at 1 month of age. To expand on this study, we could study the impact of a 2-month treatment on other ages of mice. In addition, we could also study the impact of a longer duration of treatment. A duration of two months is relatively short compared to other mouse studies using a HFD. Finally, we could also consider using different HFDs. To properly mimic a “Western Diet”, sucrose and/or cholesterol would also need to be included.

While plasma analyses are informative, future studies should expand beyond conventional blood-based methods and include imaging tools such as FibroScan, a non-invasive diagnostic tool used to assess the extent of liver fibrosis and steatosis using transient elastography, which measures the stiffness or elasticity of liver tissue.^78^ This approach is critical to the proper establishment of liver pathology.^28^ These efforts will help to establish the extent of altered BA metabolism and hepatic signaling in the population with DS. In addition, future studies should consider the impact of the gut microbiome, which plays a critical role in BA metabolism. Finally, future cohort studies will need to collect detailed information on diet from all participants to understand the contribution of diet on liver function.

## STAR⍰METHODS

### KEY RESOURCES TABLE

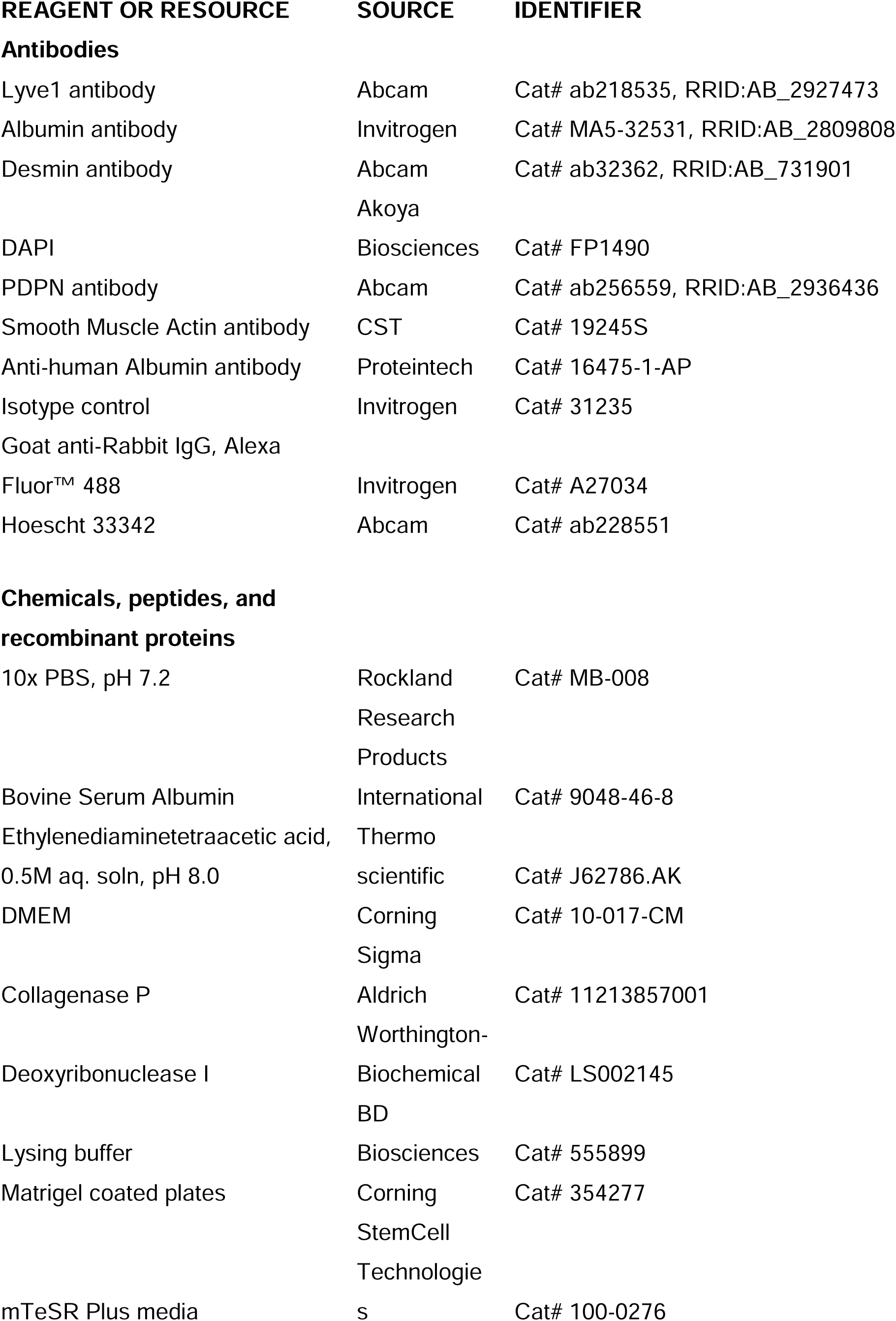
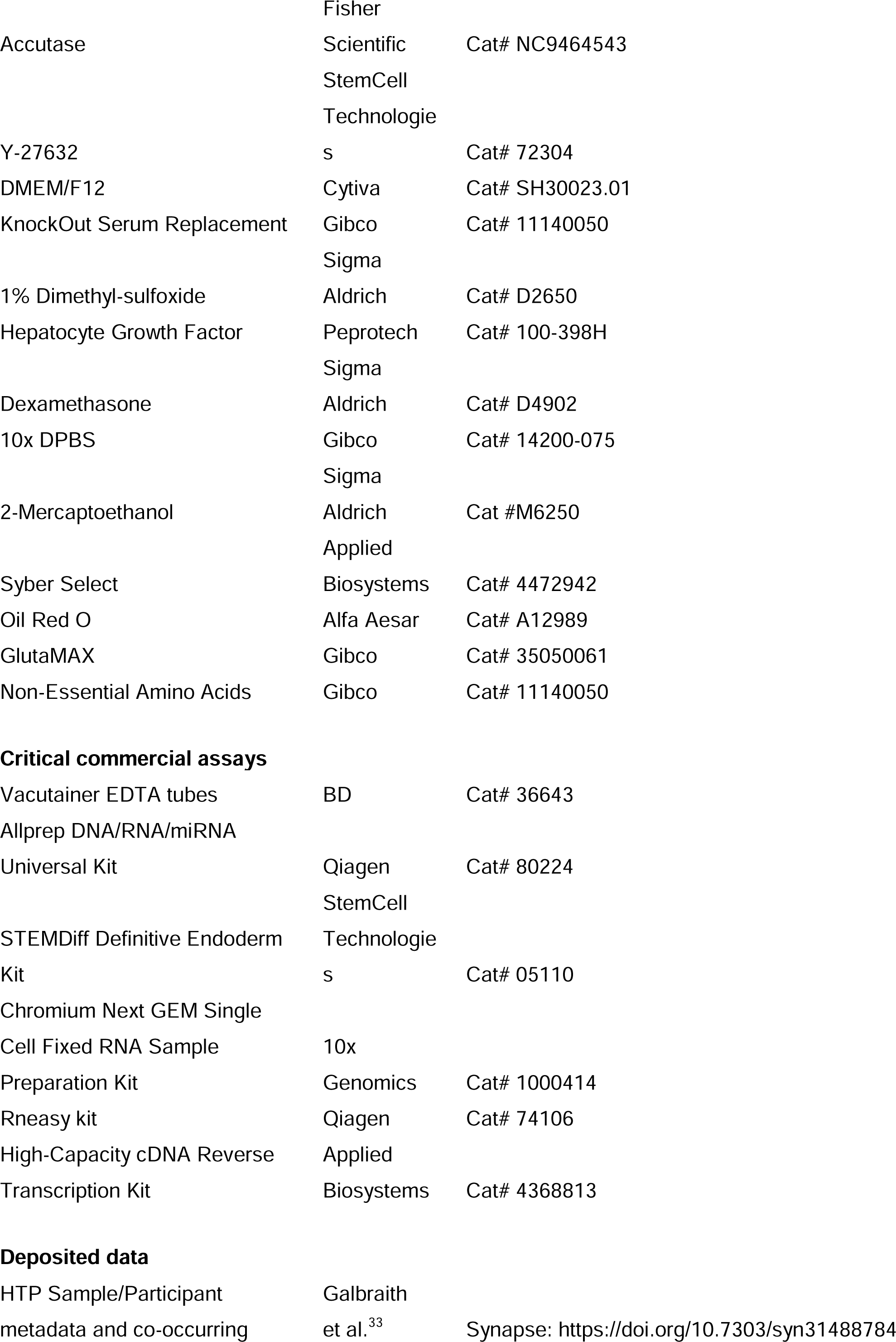
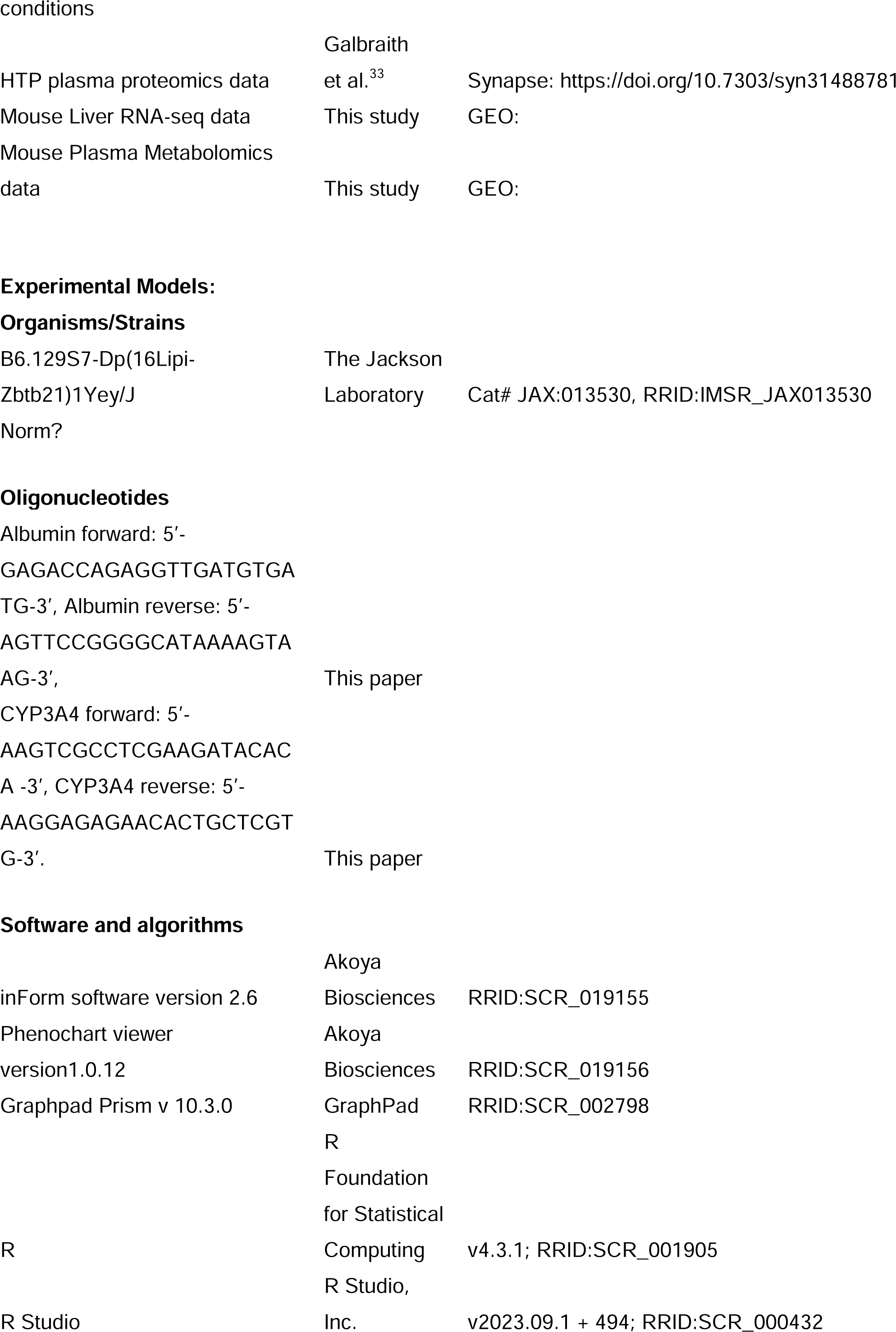
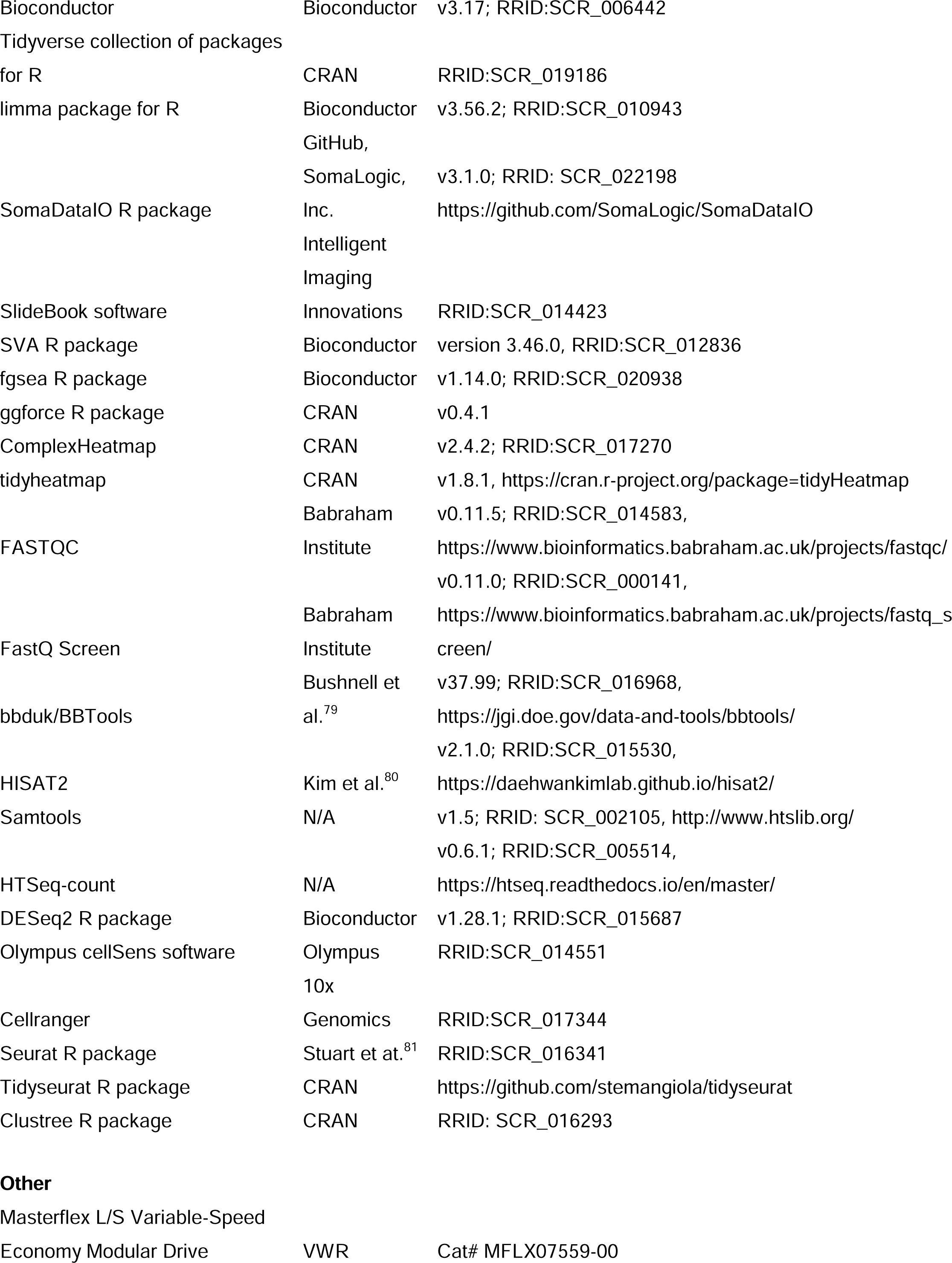
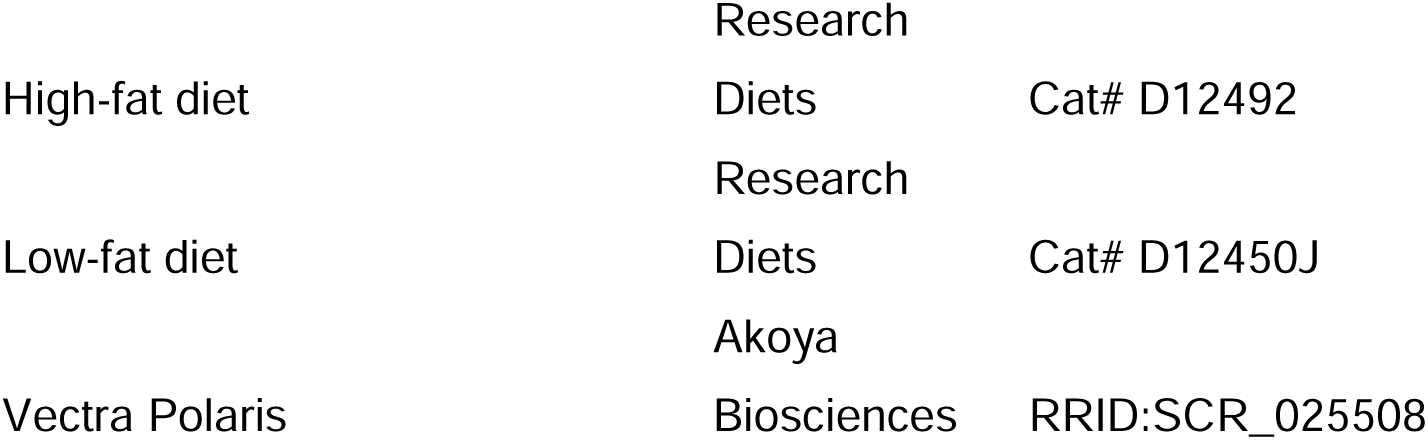

### EXPERIMENTAL MODEL AND STUDY PARTICIPANT DETAILS

#### Human Trisome Project participants

All human data in this manuscript were previously reported and reanalyzed here.^6,9^ All human study participants were enrolled in the Crnic Institute’s Human Trisome Project (HTP) under a study protocol approved by the Colorado Multiple Institutional Review Board (COMIRB 15-2170 and NCT02864108; see also www.trisome.org). Written informed consent was obtained from all study participants or their legal guardians.

#### Blood sample collection and processing

The human datasets analyzed herein were derived from peripheral blood samples collected into BD Vacutainer K2 EDTA tubes (BD, catalog no. 366643). Tubes were processed by centrifugation at 700*g* for 15 minutes to separate plasma from other blood fractions. Plasma was aliquoted, flash-frozen, and stored at −80°C. Subsequent processing was carried out as described below.

#### iPSC generation

iPSCs were generated from urine-derived renal epithelial cells or white blood cells using a cocktail of mRNAs encoding six reprogramming factors (a modified version of Oct4 fused with the MyoD transactivation domain (called M_3_O), Sox2, Klf4, cMyc, Lin28A, and Nanog) along with the reprogramming mimic miRNAs −367 and - 302 to reprogram cells.^82^

#### Hepatocyte differentiation

Hepatocyte-like cells (iHepatocytes) were differentiated from a panel of 8 iPSC lines, four control lines and four lines trisomic for chromosome 21. The iPSCs were maintained on matrigel-coated plates (Corning, Cat. # 354277) in mTeSR Plus media (StemCell Technologies, Cat. # 100-0276), manually removing any spontaneously differentiating cells. Genomic integrity and retention of the triplicated chromosome 21 in the iPSCs were validated by g-banded karyotype analysis through the University of Colorado Cancer Center Pathology Shared Resource – Cytogenetics Section.

iPSCs were differentiated into iHepatocytes using previously described protocols with minor variations^52^. Briefly, iPSCs were singularized with accutase (Fisher Scientific, Cat. # NC9464543) and seeded into matrigel-coated six-well plates at 2e^6^ cells per well in mTeSR Plus media supplemented with 10uM Y-27632 (StemCell Technologies, Cat. # 72304). After 24 hours, when cells were 90-100% confluent, Y-27632 was removed, and the iPSCs were induced to form the definitive endoderm using the STEMDiff Definitive Endoderm Kit (StemCell, Cat. # 05110) according to the manufacturer’s instructions. Day 0 of iHepatocyte induction was considered when the definitive endoderm media was placed on the cells. On day 4 of differentiation, cells were singularized and plated 7.9e^4^ cells/cm^2^ in heptoblast media supplemented with 10 uM Y-27632. Hepatoblast media consisted of DMEM/F12 (Cytiva, Cat. # SH30023.01) supplemented with 10% KnockOut Serum Replacement (KOSR, Gibco, Cat. # 10828028), 1% GlutaMAX (Gibco, Cat. # 35050061), 1% Non-Essential Amino Acids (NEAA, Gibco, Cat. # 11140050), 1% Dimethyl-sulfoxide (DMSO, Sigma-Aldrich, Cat. # D2650), and 100 ng/ml of Hepatocyte Growth Factor (HGF, Peprotech, Cat. # 100-398H). After 24 hours Y-27632 was removed and the media was changed daily until day 12. From day 12 onward, cells were cultured in hepatocyte maturation media consisting of DMEM/F12, 10% KOSR, 1% NEAA, 1% GlutaMAX, and 100 nM dexamethasone (Sigma-Aldrich, Cat. # D4902). Media was changed daily until day 15.

#### Animal husbandry and genotyping

Experiments were approved by the Institutional Animal Care and Use Committee at the University of Colorado Anschutz Medical Campus under protocol #00111 and performed in accordance with National Institutes of Health (NIH) guidelines. Dp16 (B6.129S7-Dp(16Lipi-Zbtb21)1Yey/J) mice were originally purchased from the Jackson Laboratory and maintained on the C57BL/6J background in specific pathogen–free conditions. Mice were housed separately by sex in groups of one to five mice per cage under a 14-hour light:10-hour dark cycle with controlled temperature and 35% humidity and had ad libitum access to food (16 kcal% fat diet) and water. For genotyping, genomic DNA was prepared from 1 to 2 mm of toe, tail, or ear tissue for automated genotyping by reverse transcription polymerase chain reaction with specific probes designed for *Lipi* and *Zfp295* (Transnetyx). Upon termination of the studies, all animals were euthanized by CO_2_ asphyxiation and cardiac puncture, then immediately perfused with 1x PBS using a Masterflex L/S Variable-Speed Economy Modular Drive (VWR, cat. #MFLX07559-00). Dp16^2xIfnrs^ mice were generated as described previously.^48^

### METHOD DETAILS

#### Mass spectrometry–based plasma metabolomics and lipidomics

Plasma samples were thawed on ice and extracted via a modified Folch method (chloroform/methanol/water 8:4:3). Briefly, 20 μL of sample was diluted in 130 μL of liquid chromatography–mass spectrometry grade water, 600 μL of ice-cold chloroform/methanol (2:1) was added, and the samples were vortexed for 10 seconds. Samples were then incubated at 4°C for 5 min, quickly vortexed, and centrifuged at 14,000*g* for 10 minutes at 4°C. The top (i.e., aqueous) phase was transferred to a new tube for metabolomics analysis and flash-frozen. The bottom (i.e., organic) phase was transferred to a new tube for lipidomics analysis and then dried under N_2_ flow. Analyses were performed using a Vanquish UHPLC coupled online to a Q Exactive high-resolution mass spectrometer (Thermo Fisher Scientific). Samples (10 μL per injection) were randomized and analyzed in positive and negative electrospray ionization modes (separate runs) using a 5-minute C18 gradient on a Kinetex C18 column (Phenomenex) as described.^83^ Data were analyzed using Maven in conjunction with the Kyoto Encyclopedia of Genes and Genomes database and an in-house standard library.

#### SOMAscan proteomics of human plasma

A total of 125 μL of EDTA plasma was analyzed by SOMAscan using established protocols.^6^ Each of the 4500+ SOMAmer reagents binds a target peptide and is quantified on a custom Agilent hybridization chip. Normalization and calibration were performed according to SOMAscan Data Standardization and File Specification Technical Note (SSM-020). The output of the SOMAscan assay is reported in relative fluorescence units (RFU).

#### Fibrosis/Steatosis GSEA signature

Human plasma SOMAscan proteomic data was obtained from Govaere et al, 2020.^44^ Statistical significance was defined as a Benjamini–Hochberg corrected *P-value* less than 0.05 and an absolute fold change of more than 1.25 fold. Proteins significantly more abundant in those with advanced fibrosis versus mild fibrosis, as determined by a liver biopsy and their respective NAFLD Activity Score, were used for the Fibrosis signature. The same method was used for the steatosis GSEA signature. GSEA then performed using using the HTP SOMAscan proteomic data as described below.

#### Liver Histopathology

Formalin-fixed paraffin-embedded (FFPE) pieces of liver were sectioned at 5 microns and stained with hematoxylin and eosin (H&E). Scoring of liver sections used procedures adapted for mice from the validated histological scoring system established by Kleiner et al. 2005.^46,84,85^ The pathologist was blind to animal genotype, sex, and treatment group. Histological images were captured on an Olympus BX51 microscope equipped with a 17mp Olympus DP73 high-definition color digital camera using the Olympus CellSens software (Olympus, Waltham, MA).

#### Multispectral Fluorescence Immunohistochemistry (MFI)

FFPE liver was sectioned at 5 microns, mounted on glass slides, and stained with antibodies against Albumin (hepatocytes, Invitrogen cat#MA5-32531), Lyve1 (liver endothelial cells, Abcam cat#ab218535), Desmin (activated hepatic stellate cells, Abcam cat#ab32362), PDPN (lymphatic endothelial cells, Abcam cat#ab256559), Smooth Muscle Actin (hepatic stellate cells, CST cat#19245S), and DAPI (cell nuclei, Cat# FP1490 Akoya Biosciences). A whole slide scan was performed at 20x on a Vectra Polaris instrument (Akoya Biosciences), then 6 regions of interest (ROI) per slide were selected using the Phenochart viewer version1.0.12 (Akoya Bioscience). Individual image and batch analysis was performed using inForm software version 2.6 (Akoya Biosciences). The number of cells positive for a specific cell marker as a percent of total cells per ROI was calculated and the mean per animal was used for further analysis.

#### Picrosirius red staining

FFPE liver was sectioned at 5 microns, mounted on glass slides, pre-treated with Bouin’s Fixative, and stained with Picrosirius Red. Images were obtained under polarized light in a tiling fashion and positive Cy3-42-255 pixels were quantified utilizing the SlideBook software (Intelligent Imaging Innovations, Denver, CO) and then expressed as a percentage of the total pixels examined.

#### Murine liver RNA extraction and bulk sequencing

Upon euthanasia, livers were immediately extracted and placed in 594 μL of lysis buffer RLT Plus (QIAGEN) and 6 μL of 2-mercaptoethanol (Sigma-Aldrich) for RNA extraction, then stored at −80 C. Total RNA was isolated using the AllPrep Kit (QIAGEN) following the manufacturer’s instructions. Library preparation was carried out using a Universal Plus mRNA Kit Poly(A) (Tecan). Paired-end, 150-bp sequencing was carried out on an Illumina NovaSeq 6000.

#### Analysis of bulk murine liver RNA-sequencing data

Reads were demultiplexed and converted to FASTQ format using bcl2fastq v2.20.0.422. Data quality was assessed using FASTQC v0.11.5 and FastQ Screen v0.11.0. Filtering of low-quality reads was performed using bbduk from BBTools v37.99 and fastq-mcf from ea-utils v1.05. Alignment to the mouse GRCm38 reference genome index and Gencode M24 annotation GTF was carried out using HISAT2 v2.1.0. Alignments were sorted and filtered for mapping quality (MAPQ > 10) using SAMtools v1.5. Gene-level count data were quantified using HTSeq-count v0.6.1. RNA-seq data yield was a minimum of ∼30 million raw reads. Differential gene expression was evaluated using DESeq2 v1.28.1 with surrogate variables, determined using the svaseq() function from the sva package (version 3.46.0), as covariates to remove unwanted sources of variation, including sex and batch. Significance was set at *q* < 0.1 (10% FDR). GSEA was carried out in R using the fgsea package (v 1.14.0), using Hallmark gene sets on a ranked list of log_2_-transformed fold changes.

#### Preparation of fixed single cells from mouse liver

Upon euthanasia and cardiac puncture, mouse livers were perfused with cold 1x Dulbecco’s PBS (DPBS), then removed. Livers were minced manually with a scalpel, suspended in DMEM (Corning) with 2 mg/mL Collagenase P (Roche) and 0.2 mg/mL Deoxyribonuclease I (Worthington Biochemical), and incubated with shaking at 37°C for 15 minutes. Next, the suspension was filtered through a 70 micron cell strainer to remove debris and centrifuged at 100g for 8 minutes at 4°C to separate primary mouse hepatocytes (pelleted) from the nonparenchymal cells of the liver (supernatant). Nonparenchymal cells were then pelleted by centrifugation at 400g for 7 minutes at 4°C, resuspended in red blood cell lysis buffer (BD Biosciences) and incubated for 1 minute at room temperature to remove red blood cells, filtered through a 40 micron filter, pelleted again by centrifugation at 400g for 7 minutes at 4°C, and resuspended in 50 µL DPBS with 1% (w/v) BSA (Research Products International). The hepatocyte pellet was resuspended in red blood cell lysis buffer and incubated 1 minute at room temperature, filtered through a 40 micron filter, pelleted again by centrifugation at 100g for 8 minutes at 4°C, and resuspended in 50 µL DPBS with 1% (w/v) BSA. Viability and cell counts for both hepatocyte and nonparenchymal cell fractions were determined using tryphan blue and a Countess 2 cell counter (Thermo Fisher Scientific). Per animal, each fraction was then combined at a 1:1 ratio, pelleted by centrifugation, and fixed by resuspending in a 4% (w/v) formaldehyde fixative solution from a Chromium Next GEM Single Cell Fixed RNA Sample Preparation Kit (10X Genomics).

#### Mouse liver fixed single cell library construction

Fixed liver single cell suspensions were hybridized with the Next GEM Singel Cell Fixed RNA Mouse Transcriptome Probe Kit (10X Genomics), according to manufacturer instructions, and loaded into a Chromium X (10X Genomics) microfluidics instrument to generate an oil emulsion with partitioned nanoliter-scale droplets, each ideally containing a barcoded gel bead and a single cell, along with enzyme Master Mix (10X Genomics) for probe pair ligation and gel bead primer barcode extension. The resulting Illumina-compatible sequencing library was subjected to paired-end 150 bp sequencing on a Novaseq 6000 instrument (Illumina) by the Genomics Shared Resource at the University of Colorado Anschutz Medical Campus.

#### Analysis of mouse liver single cell gene expression data

Multiplexed reads in FASTQ format were processed using Cellranger-8.0.1 in ‘multi’ mode with the mm10-2020-A transcriptome reference and Chromium Mouse Transcriptome Probe Set v1.0.1. From a total of ∼2.5 × 10^9 reads, ∼155,190 single cells were detected, with a mean of 16,141 reads per cell and 90.66% of cell barcodes passing high occupancy GEM filtering.

Cellranger-filtered per-sample barcodes, features, and matrices were read in to R as sparse matrices using the Read10X() function from the Seurat package,^86^ and per-sample Seurat objects created using the CreateSeuratObject() function, filtering to features (genes) detected in at least 3 cells (min_cells = 3) and cells with at least 200 detected features (min_features = 200). The median number cells obtained per sample was 9066 (range 6242 to 9961), with a median RNA/gene counts per cell of 4363 (range 495 to 414275), median features per cell of 2309 (range 208 to 11096), and median percent mitochondrial reads of 0.4 (range 0 to 88.6). Next, cells with detected features or mitochondrial read percentages greater than the overall median + three standard deviations were removed (N features >6398, 1.5% of total cells, and percent >6.49, 0.8% of total cells, respectively).

Normalization was carried out using the Seurat::SCTransform() function, with regression of mitochondrial percentage as a potential confounder. Integration of the SCTransform-normalized data across samples was carried out using the RPCA method of the Seurat::IntegrateLayers() function. Clustering of cells was carried out using Seurat::FindNeighbors(reduction = “integrated.rpca”, dims = 1:30) followed by Seurat::FindClusters() at a range of resolutions, and cluster cell counts and percentages per sample were obtained with the help of the tidyseurat and tidyverse/dplyr packages.^87^ Cluster stability and optimal resolution were assessed using the clustree package.^88^ For visualization, non-linear dimensional reduction was carried out using Seurat::RunUMAP(reduction = “integrated.rpca”, dims = 1:30). Qualitative color schemes were generated using the iwanthue() function from the hues package. Clusters were manually annotated with presumptive cell types based on relative expression level and percentage of cells positive for known marker genes. Changes in cluster proportion by genotype (Dp16 vs WT) were assessed using beta regression with sex as a confounding variable, and per-cluster adjustment for multiple hypothesis correction across genes using the Benjamini-Hochberg false discovery rate (FDR) method. To examine gene expression differences within cell clusters, per sample per cluster ‘pseudobulk’ gene level RNA counts were obtained using the tidyseurat:: aggregate_cells() function. Analysis of differential gene expression by genotype (Dp16 vs WT) based on pseudobulk counts was assessed using the DESeq2 package with adjustment for surrogate variables estimated using the svaseq() function from the combat package. GSEA was performed as described below.

#### Murine blood plasma analyte analysis

Blood was collected from mice via a cardiac puncture post euthanasia into lithium heparin tubes (Sarstedt). Plasma was isolated from whole blood via centrifugation for 15 minutes at 700g, then the supernatant was isolated and spun again for 15 minutes at 2100g.15 µL of plasma per analyte was processed for ALT, AST, ALP, or albumin detection using the DRI-CHEM® 7000 Veterinary Chemistry Analyzer (Heska, Loveland, Colorado) in collaboration with the Comparative Pathology Shared Resource at CU-AMC. Significance was determined using a Mann-Whitney test.

#### Q-RT-PCR

RNA was collected from samples at the iPSC (day 0), iDefinitive Endoderm (day 4), iHepatoblast (day 12), and iHepatocyte (day 15) stages using the RNeasy kit (Qiagen, Cat. #74106) according to the manufacturer’s instructions. cDNA was made using the High-Capacity cDNA Reverse Transcription Kit (Applied Biosystems, Cat. # 4368813) according to the manufacturer’s instructions. Quantitative PCR was then performed using Syber Select (Applied Biosystems, Cat. # 4472942) on the Viia7 (Applied Biosystems). Relative gene expression was determined using the delta-delta CT method normalizing to samples from the iPSC stage for Albumin or the definitive endoderm stage for CYP3A4, which was undetected in iPSCs. The following oligonucleotide sequences were used in this study. Albumin forward: 5’-GAGACCAGAGGTTGATGTGATG-3’, Albumin reverse: 5’-AGTTCCGGGGCATAAAAGTAAG-3’, CYP3A4 forward: 5’-AAGTCGCCTCGAAGATACACA −3’, CYP3A4 reverse: 5’-AAGGAGAGAACACTGCTCGTG-3’.

#### Immunofluorescence

On day 15 of differentiation, iHepatocytes were fixed in 4% paraformaldehyde (PFA) for 10 minutes at room temperature, then washed 3x for 5 minutes in PBS. Samples were permeabilized in 0.1% Triton X-100 for 10 minutes at room temperature, followed by another 3x 5 minutes series of PBS washes, and then blocked for 1 hour in a 5% bovine serum albumin solution (BSA, Research Products International, Cat. # A30075). Cells were treated with 5.4 ug/ml of either rabbit anti-human Albumin antibody (Proteintech, Cat. # 16475-1-AP) or isotype control (Invitrogen, Cat. # 31235) overnight at 4°C in 0.1% BSA and then washed 3x for 5 minutes in PBS. Samples were then incubated for 1 hour at room temperature in Goat anti-Rabbit IgG, Alexa Fluor™ 488 (Invitrogen, Cat. # A27034). Samples were washed once in PBS for 5 minutes and incubated for 10 minutes at room temperature in 5 ug/ml of Hoescht 33342 (Abcam, Cat. # ab228551). After nuclear staining, samples were washed a final time in DPBS before imaging on a Zeiss LSM 780 at the University of Colorado Anschutz Medical Campus Advanced Light Microscopy Core.

#### Oil Red O Straining

Oil Red O (ORO, Alfa Aesar, Cat. # A12989) was prepared at 3 mg/ml in 100% isopropanol. Immediately before staining, three parts of ORO were mixed with two parts of DI water, filtered through Whatman paper, and used within two hours. Day 15 iHepatocytes were fixed in 4% PFA for 10 minutes at room temperature, then washed twice in water. Cells were treated with 60% isopropanol for 5 minutes at room temperature. Isopropanol was replaced with the filtered ORO solution and incubated for 15 minutes at room temperature. After ORO staining, samples were washed with DI water until no excess stain remained. Images were captured immediately after staining using bright field microscopy.

#### High-fat diet experiments

Mice had ad libitum access to either a low-fat diet (6 kcal% fat, D12450J, Research Diets, New Brunswick, NJ) or high-fat diet (60 kcal% fat, D12492, Research Diets) for 8 weeks, beginning at 28 days of age. At the conclusion of the study, mice were euthanized as described above, tissues collected and either placed into 10% neutral buffered formalin or added to RLT/BME.

#### Statistical analyses

Data preprocessing, statistical analysis, and plot generation for all datasets were carried out using R as detailed below or Prism GraphPad. All figures were assembled in Adobe Illustrator.

#### Analysis of plasma metabolomics and lipidomics data

Metabolite relative abundance values were imported to R. Metabolites with zero values were replaced with a random value sampled from between 0 and 0.5x the minimum nonzero intensity value for that metabolite. For downstream analysis, data were then normalized using a scaling factor derived by dividing the global median intensity value across all metabolites by each sample median intensity. Extreme outliers were classified per karyotype and per analyte as measurements more than three times the interquartile range below or above the first and third quartiles, respectively, and excluded from further analysis. Differential abundance analysis for metabolites was performed using linear regression in R with log_2_-transformed relative abundance as the outcome/dependent variable; T21 status as the predictor/independent variable; and age, sex, BMI, and sample source as covariates. Multiple hypothesis correction was performed with the Benjamini-Hochberg method with an FDR threshold of 10% (*q* < 0.1). Before visualization or correlation analysis, metabolite data were adjusted for age, sex, BMI, and sample source using the removeBatchEffect() function from the limma package.

#### Analysis of SOMAscan proteomics data

Normalized data (RFU) in the SOMAscan adat file format were imported to R using the SomaDataIO R package (v3.1.0). Extreme outliers were classified per karyotype and per analyte as measurements more than three times the interquartile range (IQR) below or above the first and third quartiles, respectively (below Q1 – 3 × IQR or above Q3 + 3 × IQR), and excluded from further analysis. Differential abundance analysis for SOMAscan proteomics was performed using linear regression in R with log_2_-transformed relative abundance as the outcome/dependent variable; T21 status as the predictor/independent variable; and age, sex, BMI, and sample source as covariates. Multiple hypothesis correction was performed with the Benjamini-Hochberg method using an FDR threshold of 10% (*q* < 0.1). Before visualization or correlation analysis, SOMAscan data were adjusted for age, sex, BMI, and sample source using the removeBatchEffect() function from the limma package (v3.44.3).

#### Data visualization

For comparison of data distributions between different categories/groups, sina plots showing all points jittered horizontally by local density, modified with boxes representing medians and interquartile ranges, were generated using ggplot2 and the geom_sina() function from the ggforce R package. Heatmaps were generated using the ComplexHeatmap and tidyheatmap R packages.

#### Gene set enrichment analysis

GSEA was carried out in R using the fgsea package (v 1.14.0), using Hallmark gene sets, and log_2_-transformed fold changes (for RNA-seq), log_2_(fold change) multiplied by −log_10_(*P-value*) (for SOMAscan proteomics) as the ranking metric.

## RESOURCE AVAILABILITY

### Lead contact

Further information and requests for resources and reagents should be directed to and will be fulfilled by the lead contact, Kelly Sullivan.

### Materials availability

This study did not generate any unique materials.

### Data and code availability

All data are available in the manuscript, supplementary tables, or GEO Superseries GSE296720. Previously published plasma proteomics data using the SOMAscan platform for 410+ research participants are available through Synapse (https://doi.org/10.7303/syn31488781). Plasma metabolomics data for 410+ participants can be accessed through Synapse (https://doi.org/10.7303/syn31488782) or Metabolomics Workbench (study ID ST002200; http://dev.metabolomicsworkbench.org:22222/data/DRCCMetadata.php?Mode=Study&StudyID=S <u>T002200&Access=UrlT3545</u>).

## ACKNOWLEDGMENTS

This work was supported primarily by NIH grants R01AI145988 (K.D.S.) and R01AI150305 (J.M.E.). Additional funding was provided by NIH grants R24OD035579 (M.D.G., J.M.E., K.D.S.), P30CA046934, the Linda Crnic Institute for Down Syndrome, the Global Down Syndrome Foundation, the Anna and John J. Sie Foundation, the University of Colorado School of Medicine, and the Boettcher Foundation. We would like to thank Dr. Angelo D’Alessandro and his team at the Mass Spectrometry Metabolomics Shared Resource, Dr. Kimberly Jordan and her team at the Human Immune Monitoring Shared Resource, Dr. Schuyler (Sky) Lee and his team at the Genomics Shared Resource, the Comparative Pathology Shared Resource, especially Caterina Quartararo, and E. Erin Smith and her team at the Research Histology Shared Resource for their outstanding contributions to generating this data.

## AUTHOR CONTRIBUTIONS

LND - Investigation, visualization, methodology, writing; BFN - Investigation, visualization, methodology; NPE - Formal analysis, visualization ; KAS - Investigation; KAW - Investigation; CB - Investigation; ALR - Investigation, methodology; AET- Investigation; DJO - Investigation, methodology; MDG - Formal analysis, visualization, methodology, data curation, supervision; JME - Supervision, funding acquisition; KDS - Conceptualization, supervision, funding acquisition, visualization, writing

## DECLARATION OF GENERATIVE AI AND AI-ASSISTED TECHNOLOGIES

No generative AI was used in the preparation of this manuscript.

## SUPPLEMENTAL INFORMATION

Figs. S1 to S7

Tables S1-S9

## REFERENCES

1. Lejeune, J., Gauthier, M., and Turpin, R. (1959). [Human chromosomes in tissue cultures]. C R Hebd Seances Acad Sci 248, 602–603.

2. De Graaf, G., Buckley, F., and Skotko, B.G. (2017). Estimation of the number of people with Down syndrome in the United States. Genetics in Medicine 19, 439–447. 10.1038/gim.2016.127.

3. Antonarakis, S.E., Skotko, B.G., Rafii, M.S., Strydom, A., Pape, S.E., Bianchi, D.W., Sherman, S.L., and Reeves, R.H. (2020). Down syndrome. Nature Reviews Disease Primers 6. 10.1038/s41572-019-0143-7.

4. Sullivan, K.D., Lewis, H.C., Hill, A.A., Pandey, A., Jackson, L.P., Cabral, J.M., Smith, K.P., Liggett, L.A., Gomez, E.B., Galbraith, M.D., et al. (2016). Trisomy 21 consistently activates the interferon response. Elife 5. 10.7554/eLife.16220.

5. Araya, P., Waugh, K.A., Sullivan, K.D., Nunez, N.G., Roselli, E., Smith, K.P., Granrath, R.E., Rachubinski, A.L., Enriquez Estrada, B., Butcher, E.T., et al. (2019). Trisomy 21 dysregulates T cell lineages toward an autoimmunity-prone state associated with interferon hyperactivity. Proc Natl Acad Sci U S A 116, 24231–24241. 10.1073/pnas.1908129116.

6. Galbraith, M.D., Rachubinski, A.L., Smith, K.P., Araya, P., Waugh, K.A., Enriquez-Estrada, B., Worek, K., Granrath, R.E., Kinning, K.T., Paul Eduthan, N., et al. (2023). Multidimensional definition of the interferonopathy of Down syndrome and its response to JAK inhibition. Sci Adv 9, eadg6218. 10.1126/sciadv.adg6218.

7. Sullivan, K.D., Evans, D., Pandey, A., Hraha, T.H., Smith, K.P., Markham, N., Rachubinski, A.L., Wolter-Warmerdam, K., Hickey, F., Espinosa, J.M., and Blumenthal, T. (2017). Trisomy 21 causes changes in the circulating proteome indicative of chronic autoinflammation. Sci Rep 7, 14818. 10.1038/s41598-017-13858-3.

8. Powers, R.K., Culp-Hill, R., Ludwig, M.P., Smith, K.P., Waugh, K.A., Minter, R., Tuttle, K.D., Lewis, H.C., Rachubinski, A.L., Granrath, R.E., et al. (2019). Trisomy 21 activates the kynurenine pathway via increased dosage of interferon receptors. Nat Commun 10, 4766. 10.1038/s41467-019-12739-9.

9. Araya, P., Kinning, K.T., Coughlan, C., Smith, K.P., Granrath, R.E., Enriquez-Estrada, B.A., Worek, K., Sullivan, K.D., Rachubinski, A.L., Wolter-Warmerdam, K., et al. (2022). IGF1 deficiency integrates stunted growth and neurodegeneration in Down syndrome. Cell Rep 41, 111883. 10.1016/j.celrep.2022.111883.

10. Waugh, K.A., Araya, P., Pandey, A., Jordan, K.R., Smith, K.P., Granrath, R.E., Khanal, S., Butcher, E.T., Estrada, B.E., Rachubinski, A.L., et al. (2019). Mass Cytometry Reveals Global Immune Remodeling with Multi-lineage Hypersensitivity to Type I Interferon in Down Syndrome. Cell Rep 29, 1893–1908 e1894. 10.1016/j.celrep.2019.10.038.

11. Donovan, M.G., Eduthan, N.P., Smith, K.P., Britton, E.C., Lyford, H.R., Araya, P., Granrath, R.E., Waugh, K.A., Enriquez Estrada, B., Rachubinski, A.L., et al. (2024). Variegated overexpression of chromosome 21 genes reveals molecular and immune subtypes of Down syndrome. Nat Commun 15, 5473. 10.1038/s41467-024-49781-1.

12. Donovan, M.G., Rachubinski, A.L., Smith, K.P., Araya, P., Waugh, K.A., Enriquez-Estrada, B., Britton, E.C., Lyford, H.R., Granrath, R.E., Schade, K.A., et al. (2024). Multimodal analysis of dysregulated heme metabolism, hypoxic signaling, and stress erythropoiesis in Down syndrome. Cell Rep 43, 114599. 10.1016/j.celrep.2024.114599.

13. Lazzarino, G., Amorini, A.M., Mangione, R., Saab, M.W., Di Stasio, E., Di Rosa, M., Tavazzi, B., Lazzarino, G., Onder, G., and Carfi, A. (2022). Biochemical Discrimination of the Down Syndrome-Related Metabolic and Oxidative/Nitrosative Stress Alterations from the Physiologic Age-Related Changes through the Targeted Metabolomic Analysis of Serum. Antioxidants (Basel) 11. 10.3390/antiox11061208.

14. Anderson, C.C., Marentette, J.O., Prutton, K.M., Rauniyar, A.K., Reisz, J.A., D’Alessandro, A., Maclean, K.N., Saba, L.M., and Roede, J.R. (2021). Trisomy 21 results in modest impacts on mitochondrial function and central carbon metabolism. Free Radic Biol Med 172, 201–212. 10.1016/j.freeradbiomed.2021.06.003.

15. Antonaros, F., Ghini, V., Pulina, F., Ramacieri, G., Cicchini, E., Mannini, E., Martelli, A., Feliciello, A., Lanfranchi, S., Onnivello, S., et al. (2020). Plasma metabolome and cognitive skills in Down syndrome. Sci Rep 10, 10491. 10.1038/s41598-020-67195-z.

16. Culp-Hill, R., Zheng, C., Reisz, J.A., Smith, K., Rachubinski, A., Nemkov, T., Butcher, E., Granrath, R., Hansen, K.C., Espinosa, J.M., and D’Alessandro, A. (2017). Red blood cell metabolism in Down syndrome: hints on metabolic derangements in aging. Blood Adv 1, 2776–2780. 10.1182/bloodadvances.2017011957.

17. Oreskovic, N.M., Baumer, N.T., Di Camillo, C., Cornachia, M., Franklin, C., Hart, S.J., Kishnani, P.S., McCormick, A., Milliken, A.L., Patsiogiannis, V., et al. (2023). Cardiometabolic profiles in children and adults with overweight and obesity and down syndrome. Am J Med Genet A 191, 813–822. 10.1002/ajmg.a.63088.

18. Fleishman, J.S., and Kumar, S. (2024). Bile acid metabolism and signaling in health and disease: molecular mechanisms and therapeutic targets. Signal Transduction and Targeted Therapy 9, 97. 10.1038/s41392-024-01811-6.

19. Li, X.J., Fang, C., Zhao, R.H., Zou, L., Miao, H., and Zhao, Y.Y. (2024). Bile acid metabolism in health and ageing-related diseases. Biochem Pharmacol 225, 116313. 10.1016/j.bcp.2024.116313.

20. Long, J., Xu, Y., Zhang, X., Wu, B., and Wang, C. (2024). Role of FXR in the development of NAFLD and intervention strategies of small molecules. Arch Biochem Biophys 757, 110024. 10.1016/j.abb.2024.110024.

21. Jin, W., Zheng, M., Chen, Y., and Xiong, H. (2024). Update on the development of TGR5 agonists for human diseases. Eur J Med Chem 271, 116462. 10.1016/j.ejmech.2024.116462.

22. Fuchs, C.D., Simbrunner, B., Baumgartner, M., Campbell, C., Reiberger, T., and Trauner, M. (2025). Bile acid metabolism and signalling in liver disease. Journal of Hepatology 82, 134–153. 10.1016/j.jhep.2024.09.032.

23. Foundation, A.L. (2023). How Many People Have Liver Disease? https://liverfoundation.org/about-your-liver/facts-about-liver-disease/how-many-people-have-liver-disease/.

24. Rinella, M.E., Lazarus, J.V., Ratziu, V., Francque, S.M., Sanyal, A.J., Kanwal, F., Romero, D., Abdelmalek, M.F., Anstee, Q.M., Arab, J.P., et al. (2023). A multi-society Delphi consensus statement on new fatty liver disease nomenclature. Ann Hepatol, 101133. 10.1016/j.aohep.2023.101133.

25. Huang, S., Bao, Y., Zhang, N., Niu, R., and Tian, L. (2023). Long-term outcomes in lean and non-lean NAFLD patients: a systematic review and meta-analysis. Endocrine. 10.1007/s12020-023-03351-5.

26. Dufour, D.R., Lott, J.A., Nolte, F.S., Gretch, D.R., Koff, R.S., and Seeff, L.B. (2000). Diagnosis and monitoring of hepatic injury. II. Recommendations for use of laboratory tests in screening, diagnosis, and monitoring. Clin Chem 46, 2050–2068. 10.1093/clinchem/46.12.2050.

27. Jayasekera, D., and Hartmann, P. (2023). Noninvasive biomarkers in pediatric nonalcoholic fatty liver disease. World J Hepatol 15, 609–640. 10.4254/wjh.v15.i5.609.

28. Jang, W., and Song, J.S. (2023). Non-Invasive Imaging Methods to Evaluate Non-Alcoholic Fatty Liver Disease with Fat Quantification: A Review. Diagnostics (Basel) 13. 10.3390/diagnostics13111852.

29. Baksh, R.A., Pape, S.E., Chan, L.F., Aslam, A.A., Gulliford, M.C., Strydom, A., and Consortium, G.-D. (2023). Multiple morbidity across the lifespan in people with Down syndrome or intellectual disabilities: a population-based cohort study using electronic health records. Lancet Public Health 8, e453–e462. 10.1016/S2468-2667(23)00057-9.

30. Valentini, D., Alisi, A., di Camillo, C., Sartorelli, M.R., Crudele, A., Bartuli, A., Nobili, V., and Villani, A. (2017). Nonalcoholic Fatty Liver Disease in Italian Children with Down Syndrome: Prevalence and Correlation with Obesity-Related Features. J Pediatr 189, 92–97 e91. 10.1016/j.jpeds.2017.05.077.

31. Tuttle, K.D., Waugh, K.A., Araya, P., Minter, R., Orlicky, D.J., Ludwig, M., Andrysik, Z., Burchill, M.A., Tamburini, B.A.J., Sempeck, C., et al. (2020). JAK1 Inhibition Blocks Lethal Immune Hypersensitivity in a Mouse Model of Down Syndrome. Cell Rep 33, 108407. 10.1016/j.celrep.2020.108407.

32. Donovan, M.G., Rachubinski, A.L., Smith, K.P., Araya, P., Waugh, K.A., Enriquez-Estrada, B., Britton, E.C., Lyford, H.R., Granrath, R.E., Schade, K.A., et al. (2024). Multimodal analysis of dysregulated heme metabolism, hypoxic signaling, and stress erythropoiesis in Down syndrome. Cell Reports 43. 10.1016/j.celrep.2024.114599.

33. Galbraith, M.D., Rachubinski, A.L., Smith, K.P., Araya, P., Waugh, K.A., Enriquez-Estrada, B., Worek, K., Granrath, R.E., Kinning, K.T., Paul Eduthan, N., et al. (2023). Multidimensional definition of the interferonopathy of Down syndrome and its response to JAK inhibition. Science Advances 9. 10.1126/sciadv.adg6218.

34. Araya, P., Kinning, K.T., Coughlan, C., Smith, K.P., Granrath, R.E., Enriquez-Estrada, B.A., Worek, K., Sullivan, K.D., Rachubinski, A.L., Wolter-Warmerdam, K., et al. (2022). IGF1 deficiency integrates stunted growth and neurodegeneration in Down syndrome. Cell Reports 41, 111883. 10.1016/j.celrep.2022.111883.

35. Rachubinski, A.L., Wallace, E., Gurnee, E., Enriquez Estrada, B.A., Worek, K.R., Smith, K.P., Araya, P., Waugh, K.A., Granrath, R.E., Britton, E., et al. (2024). JAK inhibition decreases the autoimmune burden in Down syndrome. eLife Sciences Publications, Ltd.

36. Yu, T., Liu, C., Belichenko, P., Clapcote, S.J., Li, S., Pao, A., Kleschevnikov, A., Bechard, A.R., Asrar, S., Chen, R., et al. (2010). Effects of individual segmental trisomies of human chromosome 21 syntenic regions on hippocampal long-term potentiation and cognitive behaviors in mice. Brain Res 1366, 162–171. 10.1016/j.brainres.2010.09.107.

37. Gold, L., Walker, J.J., Wilcox, S.K., and Williams, S. (2012). Advances in human proteomics at high scale with the SOMAscan proteomics platform. New biotechnology. 29, 543–549. 10.1016/j.nbt.2011.11.016.

38. Anneren, G., Gustavson, K.H., Sara, V.R., and Tuvemo, T. (1990). Growth retardation in Down syndrome in relation to insulin-like growth factors and growth hormone. Am J Med Genet Suppl 7, 59–62. 10.1002/ajmg.1320370710.

39. Luo, Y., Wadhawan, S., Greenfield, A., Decato, B.E., Oseini, A.M., Collen, R., Shevell, D.E., Thompson, J., Jarai, G., Charles, E.D., and Sanyal, A.J. (2021). SOMAscan Proteomics Identifies Serum Biomarkers Associated With Liver Fibrosis in Patients With NASH. Hepatol Commun 5, 760–773. 10.1002/hep4.1670.

40. Li, G., Shen, Q., Xu, H., Zhou, Y., Li, C., Li, Y., and He, M. (2023). SAA1 identified as a potential prediction biomarker for metastasis of hepatocellular carcinoma via multi-omics approaches. Frontiers in Oncology 13. 10.3389/fonc.2023.1138995.

41. Kerbert, A.J.C., Gupta, S., Alabsawy, E., Dobler, I., Lønsmann, I., Hall, A., Nielsen, S.H., Nielsen, M.J., Gronbaek, H., Amoros, À., et al. (2021). Biomarkers of extracellular matrix formation are associated with acute-on-chronic liver failure. JHEP Rep 3, 100355. 10.1016/j.jhepr.2021.100355.

42. Hribal, M.L., Procopio, T., Petta, S., Sciacqua, A., Grimaudo, S., Pipitone, R.M., Perticone, F., and Sesti, G. (2013). Insulin-Like Growth Factor-I, Inflammatory Proteins, and Fibrosis in Subjects With Nonalcoholic Fatty Liver Disease. The Journal of Clinical Endocrinology & Metabolism 98, E304–E308. 10.1210/jc.2012-3290.

43. Ichikawa, T., Nakao, K., Hamasaki, K., Furukawa, R., Tsuruta, S., Ueda, Y., Taura, N., Shibata, H., Fujimoto, M., Toriyama, K., and Eguchi, K. (2007). Role of growth hormone, insulin-like growth factor 1 and insulin-like growth factor-binding protein 3 in development of non-alcoholic fatty liver disease. Hepatology International 1, 287–294. 10.1007/s12072-007-9007-4.

44. Govaere, O., Hasoon, M., Alexander, L., Cockell, S., Tiniakos, D., Ekstedt, M., Schattenberg, J.M., Boursier, J., Bugianesi, E., Ratziu, V., et al. (2023). A proteo-transcriptomic map of non-alcoholic fatty liver disease signatures. Nature Metabolism 5, 572–578. 10.1038/s42255-023-00775-1.

45. Sarver, D.C., Xu, C., Velez, L.M., Aja, S., Jaffe, A.E., Seldin, M.M., Reeves, R.H., and Wong, G.W. (2023). Dysregulated systemic metabolism in a Down syndrome mouse model. Molecular Metabolism 68, 101666. 10.1016/j.molmet.2022.101666.

46. Tuttle, K.D., Waugh, K.A., Araya, P., Minter, R., Orlicky, D.J., Ludwig, M., Andrysik, Z., Burchill, M.A., Tamburini, B.A.J., Sempeck, C., et al. (2020). JAK1 Inhibition Blocks Lethal Immune Hypersensitivity in a Mouse Model of Down Syndrome. Cell Reports 33, 108407. 10.1016/j.celrep.2020.108407.

47. Kisseleva, T., and Brenner, D. (2021). Molecular and cellular mechanisms of liver fibrosis and its regression. Nature Reviews Gastroenterology & Hepatology 18, 151–166. 10.1038/s41575-020-00372-7.

48. Waugh, K.A., Minter, R., Baxter, J., Chi, C., Galbraith, M.D., Tuttle, K.D., Eduthan, N.P., Kinning, K.T., Andrysik, Z., Araya, P., et al. (2023). Triplication of the interferon receptor locus contributes to hallmarks of Down syndrome in a mouse model. Nature Genetics 55, 1034–1047. 10.1038/s41588-023-01399-7.

49. Gazzinelli, R.T., Wysocka, M., Hieny, S., Scharton-Kersten, T., Cheever, A., Kühn, R., Müller, W., Trinchieri, G., and Sher, A. (1996). In the absence of endogenous IL-10, mice acutely infected with Toxoplasma gondii succumb to a lethal immune response dependent on CD4+ T cells and accompanied by overproduction of IL-12, IFN-gamma and TNF-alpha. J Immunol 157, 798–805.

50. Iyer, S.S., and Cheng, G. (2012). Role of interleukin 10 transcriptional regulation in inflammation and autoimmune disease. Crit Rev Immunol 32, 23–63. 10.1615/critrevimmunol.v32.i1.30.

51. Claudel, T., Sturm, E., Duez, H., Torra, I.P., Sirvent, A., Kosykh, V., Fruchart, J.C., Dallongeville, J., Hum, D.W., Kuipers, F., and Staels, B. (2002). Bile acid-activated nuclear receptor FXR suppresses apolipoprotein A-I transcription via a negative FXR response element. J Clin Invest 109, 961–971. 10.1172/JCI14505.

52. Carpentier, A., Nimgaonkar, I., Chu, V., Xia, Y., Hu, Z., and Liang, T.J. (2016). Hepatic differentiation of human pluripotent stem cells in miniaturized format suitable for high-throughput screen. Stem Cell Res 16, 640–650. 10.1016/j.scr.2016.03.009.

53. Ahmadi, A., Niknahad, H., Li, H., Mobasheri, A., Manthari, R.K., Azarpira, N., Mousavi, K., Khalvati, B., Zhao, Y., Sun, J., et al. (2021). The inhibition of NFsmall ka, CyrillicB signaling and inflammatory response as a strategy for blunting bile acid-induced hepatic and renal toxicity. Toxicol Lett 349, 12–29. 10.1016/j.toxlet.2021.05.012.

54. Heidari, R., Jamshidzadeh, A., Ommati, M.M., Rashidi, E., Khodaei, F., Sadeghi, A., Hosseini, A., and Niknahad, H. (2019). Ammonia-induced mitochondrial impairment is intensified by manganese co-exposure: relevance to the management of subclinical hepatic encephalopathy and cirrhosis-associated brain injury. Clin Exp Hepatol 5, 109–117. 10.5114/ceh.2019.85071.

55. Ommati, M.M., Sabouri, S., Niknahad, H., Arjmand, A., Alidaee, S., Mazloomi, S., Najibi, A., Rezaei, H., Ghiasvand, A., Ahmadi, P., et al. (2023). Pulmonary inflammation, oxidative stress, and fibrosis in a mouse model of cholestasis: the potential protective properties of the dipeptide carnosine. Naunyn Schmiedebergs Arch Pharmacol 396, 1129–1142. 10.1007/s00210-023-02391-y.

56. Ommati, M.M., Farshad, O., Niknahad, H., Arabnezhad, M.R., Azarpira, N., Mohammadi, H.R., Haghnegahdar, M., Mousavi, K., Akrami, S., Jamshidzadeh, A., and Heidari, R. (2019). Cholestasis-associated reproductive toxicity in male and female rats: The fundamental role of mitochondrial impairment and oxidative stress. Toxicol Lett 316, 60–72. 10.1016/j.toxlet.2019.09.009.

57. Ptomey, L.T., Oreskovic, N.M., Hendrix, J.A., Nichols, D., and Agiovlasitis, S. (2022). Weight management recommendations for youth with Down syndrome: Expert recommendations. Front Pediatr 10, 1064108. 10.3389/fped.2022.1064108.

58. Calcaterra, V., Gazzarri, A., De Silvestri, A., Madia, C., Baldassarre, P., Rossi, V., Garella, V., and Zuccotti, G. (2023). Thyroid function, sensitivity to thyroid hormones, and metabolic syndrome in euthyroid children and adolescents with Down syndrome. J Endocrinol Invest. 10.1007/s40618-023-02086-4.

59. Ismaiel, A., Hosiny, B.E., Ismaiel, M., Leucuta, D.C., Popa, S.L., Catana, C.S., and Dumitrascu, D.L. (2023). Waist to Height Ratio in Nonalcoholic Fatty Liver Disease - Systematic Review and Meta-analysis. Clin Res Hepatol Gastroenterol, 102160. 10.1016/j.clinre.2023.102160.

60. Reiter, F.P., Hadjamu, N.J., Nagdyman, N., Zachoval, R., Mayerle, J., De Toni, E.N., Kaemmerer, H., and Denk, G. (2021). Congenital heart disease-associated liver disease: a narrative review. Cardiovasc Diagn Ther 11, 577–590. 10.21037/cdt-20-595.

61. Narciso-Schiavon, J.L., and Schiavon, L.L. (2023). Fatty liver and celiac disease: Why worry? World J Hepatol 15, 666–674. 10.4254/wjh.v15.i5.666.

62. Janota, B., Szczepanska, E., Adamek, B., and Janczewska, E. (2023). Hypothyroidism and non-alcoholic fatty liver disease: A coincidence or a causal relationship? World J Hepatol 15, 641–648. 10.4254/wjh.v15.i5.641.

63. Tehranchinia, Z., Abdollahimajd, F., Haghighatkhah, H., Talebi, A., Yarahmadi, A., and Zoghi, G. (2023). The frequency of fatty liver in patients with alopecia areata: A case-control study. J Cosmet Dermatol. 10.1111/jocd.15754.

64. Terziroli Beretta-Piccoli, B., Invernizzi, P., Gershwin, M.E., and Mainetti, C. (2017). Skin Manifestations Associated with Autoimmune Liver Diseases: a Systematic Review. Clin Rev Allergy Immunol 53, 394–412. 10.1007/s12016-017-8649-9.

65. Kelty, T.J., Dashek, R.J., Arnold, W.D., and Rector, R.S. (2023). Emerging Links between Nonalcoholic Fatty Liver Disease and Neurodegeneration. Semin Liver Dis 43, 77–88. 10.1055/s-0043-1762585.

66. Adelekan, T., Magge, S., Shults, J., Stallings, V., and Stettler, N. (2012). Lipid profiles of children with Down syndrome compared with their siblings. Pediatrics 129, e1382–e1387.

67. Ordóñez-Munoz, F.J., Rosety-Rodríguez, M., Rosety-Rodríguez, J.M., and Rosety-Plaza, M. (2005). Anthropometrical measurements as predictor of serum lipid profile in adolescents with Down syndrome. Revista de investigación clínica 57, 691–694.

68. Garcia-de la Puente, S., Flores-Arizmendi, K.A., Delgado-Montemayor, M.J., and Vargas-Robledo, T.T. (2021). Lipid profile of Mexican children with Down syndrome. BMC Pediatr 21, 77. 10.1186/s12887-021-02542-1.

69. de la Piedra, M.J., Alberti, G., Cerda, J., Cárdenas, A., Paul, M.A., and Lizama, M. (2017). [High frequency of dyslipidemia in children and adolescents with Down Syndrome]. Rev Chil Pediatr 88, 595–601. 10.4067/s0370-41062017000500004.

70. Fiorucci, S., Biagioli, M., Zampella, A., and Distrutti, E. (2018). Bile Acids Activated Receptors Regulate Innate Immunity. Front Immunol 9, 1853. 10.3389/fimmu.2018.01853.

71. Lefebvre, P., Cariou, B., Lien, F., Kuipers, F., and Staels, B. (2009). Role of bile acids and bile acid receptors in metabolic regulation. Physiol Rev 89, 147–191. 10.1152/physrev.00010.2008.

72. Evangelakos, I., Heeren, J., Verkade, E., and Kuipers, F. (2021). Role of bile acids in inflammatory liver diseases. Semin Immunopathol 43, 577–590. 10.1007/s00281-021-00869-6.

73. Kamimoto, K., Nakano, Y., Kaneko, K., Miyajima, A., and Itoh, T. (2020). Multidimensional imaging of liver injury repair in mice reveals fundamental role of the ductular reaction. Communications Biology 3, 289. 10.1038/s42003-020-1006-1.

74. Rui, L. (2014). Energy metabolism in the liver. Compr Physiol 4, 177–197. 10.1002/cphy.c130024.

75. Demetrius, L. (2005). Of mice and men. EMBO reports 6, S39–S44. 10.1038/sj.embor.7400422.

76. Perlman, R.L. (2016). Mouse Models of Human Disease: An Evolutionary Perspective. Evolution, Medicine, and Public Health, eow014. 10.1093/emph/eow014.

77. Li, Z., Yu, T., Morishima, M., Pao, A., LaDuca, J., Conroy, J., Nowak, N., Matsui, S.-I., Shiraishi, I., and Yu, Y.E. (2007). Duplication of the entire 22.9 Mb human chromosome 21 syntenic region on mouse chromosome 16 causes cardiovascular and gastrointestinal abnormalities. Human Molecular Genetics 16, 1359–1366. 10.1093/hmg/ddm086.

78. Afdhal, N.H. (2012). Fibroscan (transient elastography) for the measurement of liver fibrosis. Gastroenterol Hepatol (N Y) 8, 605–607.

79. Bushnell, B., Rood, J., and Singer, E. (2017). BBMerge – Accurate paired shotgun read merging via overlap. PLOS ONE 12, e0185056. 10.1371/journal.pone.0185056.

80. Kim, D., Paggi, J.M., Park, C., Bennett, C., and Salzberg, S.L. (2019). Graph-based genome alignment and genotyping with HISAT2 and HISAT-genotype. Nature Biotechnology 37, 907–915. 10.1038/s41587-019-0201-4.

81. Stuart, T., Butler, A., Hoffman, P., Hafemeister, C., Papalexi, E., Mauck, W.M., Hao, Y., Stoeckius, M., Smibert, P., and Satija, R. (2019). Comprehensive Integration of Single-Cell Data. Cell 177, 1888–1902.e1821. 10.1016/j.cell.2019.05.031.

82. Kogut, I., McCarthy, S.M., Pavlova, M., Astling, D.P., Chen, X., Jakimenko, A., Jones, K.L., Getahun, A., Cambier, J.C., Pasmooij, A.M.G., et al. (2018). High-efficiency RNA-based reprogramming of human primary fibroblasts. Nat Commun 9, 745. 10.1038/s41467-018-03190-3.

83. Nemkov, T., Reisz, J.A., Gehrke, S., Hansen, K.C., and D’Alessandro, A. (2019). High-Throughput Metabolomics: Isocratic and Gradient Mass Spectrometry-Based Methods. In High-Throughput Metabolomics: Methods and Protocols, A. D’Alessandro, ed. (Springer New York), pp. 13–26. 10.1007/978-1-4939-9236-2_2.

84. Lanaspa, M.A., Andres-Hernando, A., Orlicky, D.J., Cicerchi, C., Jang, C., Li, N., Milagres, T., Kuwabara, M., Wempe, M.F., Rabinowitz, J.D., et al. (2018). Ketohexokinase C blockade ameliorates fructose-induced metabolic dysfunction in fructose-sensitive mice. J Clin Invest 128, 2226–2238. 10.1172/JCI94427.

85. Kleiner, D.E., Brunt, E.M., Van Natta, M., Behling, C., Contos, M.J., Cummings, O.W., Ferrell, L.D., Liu, Y.C., Torbenson, M.S., Unalp-Arida, A., et al. (2005). Design and validation of a histological scoring system for nonalcoholic fatty liver disease. Hepatology 41, 1313–1321. 10.1002/hep.20701.

86. Satija, R., Farrell, J.A., Gennert, D., Schier, A.F., and Regev, A. (2015). Spatial reconstruction of single-cell gene expression data. Nat Biotechnol 33, 495–502. 10.1038/nbt.3192.

87. Mangiola, S., Doyle, M.A., and Papenfuss, A.T. (2021). Interfacing Seurat with the R tidy universe. Bioinformatics 37, 4100–4107. 10.1093/bioinformatics/btab404.

88. Zappia, L., and Oshlack, A. (2018). Clustering trees: a visualization for evaluating clusterings at multiple resolutions. Gigascience 7. 10.1093/gigascience/giy083.

