## Supplementary Figures and Legends for "Altered Hepatic Metabolism in Down Syndrome"

***Dunn et al., Supplementary Figure 1***

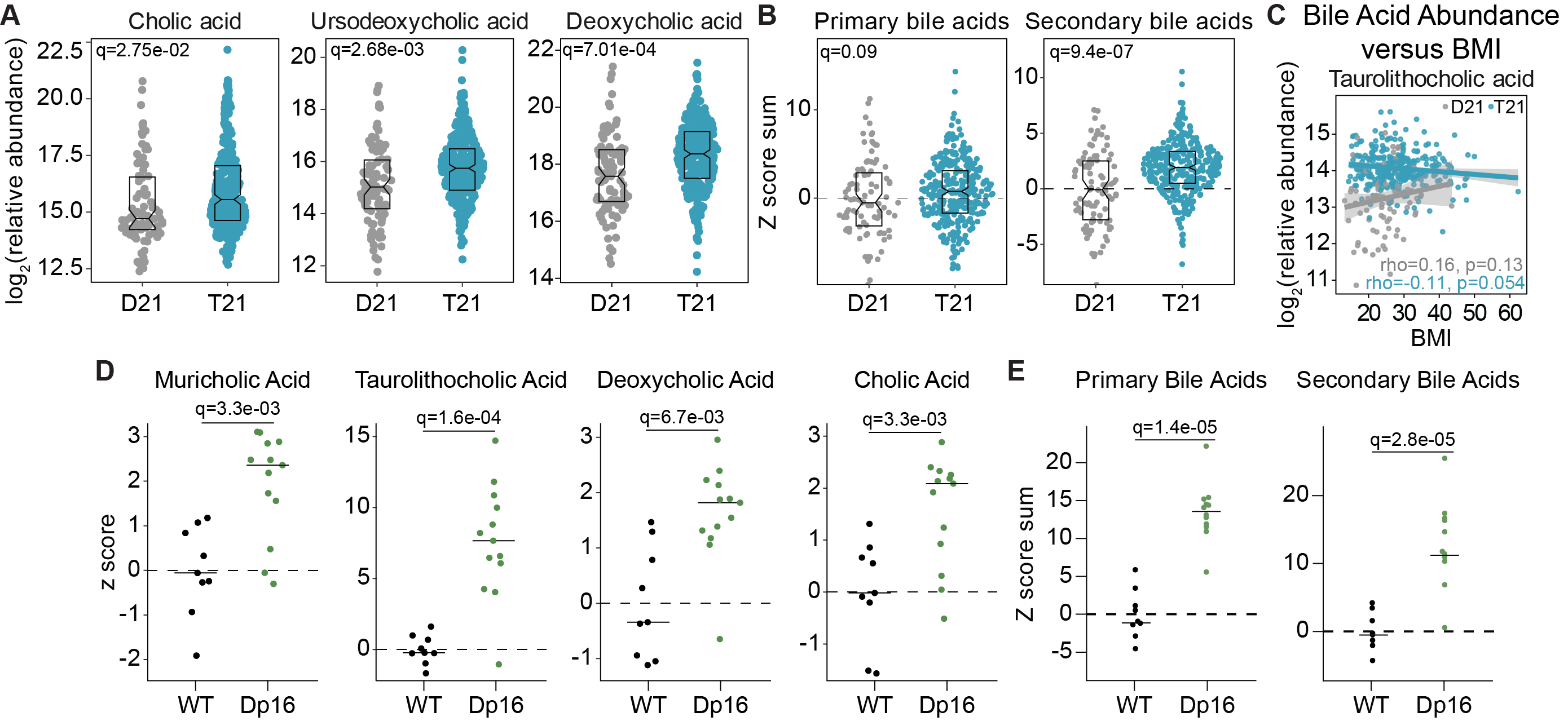

**Supplementary Figure 1. Altered bile acid metabolism in Down syndrome.**

(**A**) Sina plots displaying the log_2_(relative abundance) of cholic acid and ursodeoxycholic acid plasma metabolites in disomic individuals and those with DS. Benjamini-Hochberg adjusted p-values (q-values) are indicated.

(**B**) Sina plots displaying the sum of the z scores of both primary and secondary bile acids from disomic individuals and those with DS. Benjamini-Hochberg adjusted p-values (q-values) are indicated.

(**C**) Scatter plot displaying the relationship between relative abundance of taurolithocholic acid and BMI both for D21 individuals and those with DS. Spearman rho values and p-values are shown.

(**D**) Sina plots displaying the z score of muricholic acid, taurolithocholic acid, deoxycholic acid, and cholic acid plasma metabolites from WT and Dp16 mice. Benjamini-Hochberg adjusted p-values (q-values) are indicated.

(**E**) Sina plots displaying the sum of the z scores of both primary and secondary bile acids from WT and Dp16 mice.

Boxes in Sina plots represent interquartile ranges and medians, with notches approximating 95% confidence intervals. Benjamini-Hochberg adjusted p-values (q-values) are indicated.

***Dunn et al., Supplementary Figure 2***

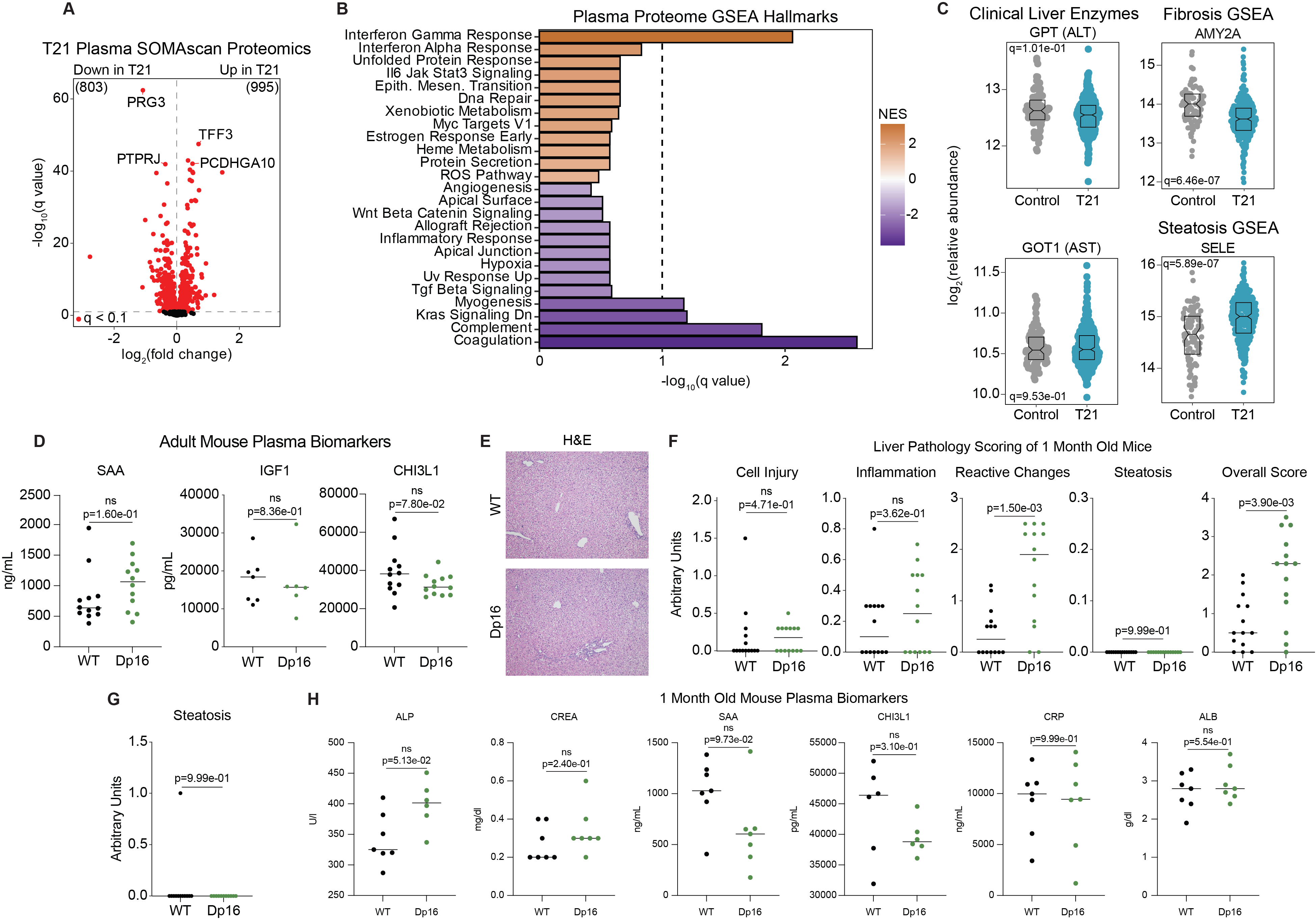

**Supplementary Figure 2. Evidence for widespread liver dysfunction in Down syndrome.**

(**A**) Volcano plot summarizing results of plasma SOMAscan proteomic analysis of individuals with DS versus disomic individuals. Significantly differentially abundant (q<0.1) proteins are colored in red.

(**B**) Bar plot summarizing results of GSEA of plasma SOMAscan proteomic analysis from individuals with DS versus disomic individuals. Dotted line indicates significance threshold of q-value less than 0.1.

(**C**) Sina plots displaying the log_2_(relative abundance) of various plasma proteins in disomic individuals and those with DS. Individual data are presented with a bar at the median and p-values, as determined by Mann-Whitney U test, are shown.

(**D**) Sina plots displaying the concentration of various plasma proteins in adult WT (n=7-12, 3-6 females) and Dp16 (n=6-12, 3-6 females) mice. Individual data are presented with a bar at the median and p-values, as determined by Mann-Whitney U test, are shown.

(**E**) Representative images of H&E-stained liver sections from one month old wildtype and Dp16 mice.

(**F**) Sina plots showing the metrics of liver pathology scoring from one month old wildtype (n=14, 8 females) and Dp16 (n=14, 6 females) mice. Individual data are presented with a bar at the median and P- values have been determined by Mann-Whitney U test.

(**G**) Sina plot displaying steatosis liver pathology scores for adult WT and Dp16 mice. P- value has been determined by Mann-Whitney U test.

(**H**) Sina plots displaying the concentration of various plasma proteins in one month old WT (n=6-7,3-5 females) and Dp16 (n=6-7,3-4 females) mice. Individual data are presented with a bar at the median and p-values, as determined by Mann-Whitney U test, are shown.

***Dunn et al., Supplementary Figure 3***

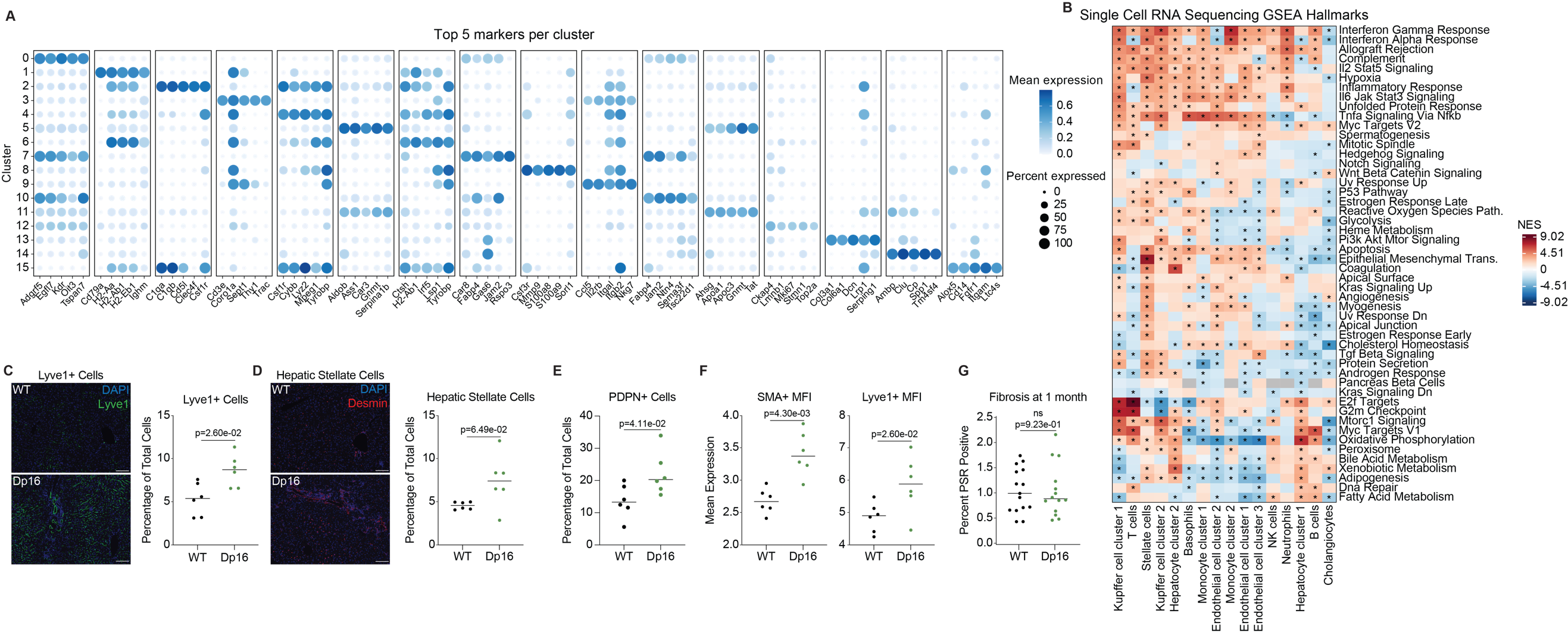

**Supplementary** **Figure 3. Stellate cell dysfunction drives liver fibrosis in the Dp16 model.**

(**A**) Bubble plot displaying the top 5 markers per cluster, with color representing mean expression and size representing percent of cells with expression.

(**B**) Heatmap displaying the enrichment scores of various GSEA pathways in each cluster from single cell RNA sequencing analysis. Asterisks denote significance (q<0.1). Color coding indicates a positive (red) or a negative (blue) normalized enrichment score (NES).

(**C**) Representative images of multiplexed stained liver sections from wildtype and Dp16 mice where DAPI is stained in blue and Lyve1 in green. Sina plot shows number of Lyve1+ cells as a percent of total cells for wildtype mice (n=6, 3 females) and Dp16 mice (n=6, 3 females). Individual data are presented with a bar at the median and the p-value, as determined by Mann-Whitney U test, is shown.

(**D**) Representative images of multiplexed stained liver sections from wildtype and Dp16 mice where DAPI is stained in blue and desmin in red. Sina plot shows number of desmin+ cells as a percent of total cells for wildtype mice (n=6, 3 females) and Dp16 mice (n=6, 3 females). Individual data are presented with a bar at the median and the p-value, as determined by Mann-Whitney U test, is shown.

(**E**) Sina plot shows number of PDPN+ cells as a percent of total cells for wildtype mice (n=6, 3 females) and Dp16 mice (n=6, 3 females). Individual data are presented with a bar at the median and the p-value, as determined by Mann-Whitney U test, is shown.

(**F**) Sina plots show mean expression of smooth muscle actin (SMA) and Lyve1 per animal for both wildtype mice (n=6, 3 females) and Dp16 mice (n=6, 3 females). Individual data are presented with a bar at the median and the p-value, as determined by Mann-Whitney U test, is shown.

(**G**) Sina plot shows the percent PSR positive pixels in wildtype and Dp16 mice at one month of age. Individual data are presented with a bar at the median and the p-value, as determined by Mann-Whitney U test, is shown.

***Dunn et al., Supplementary Figure 4***

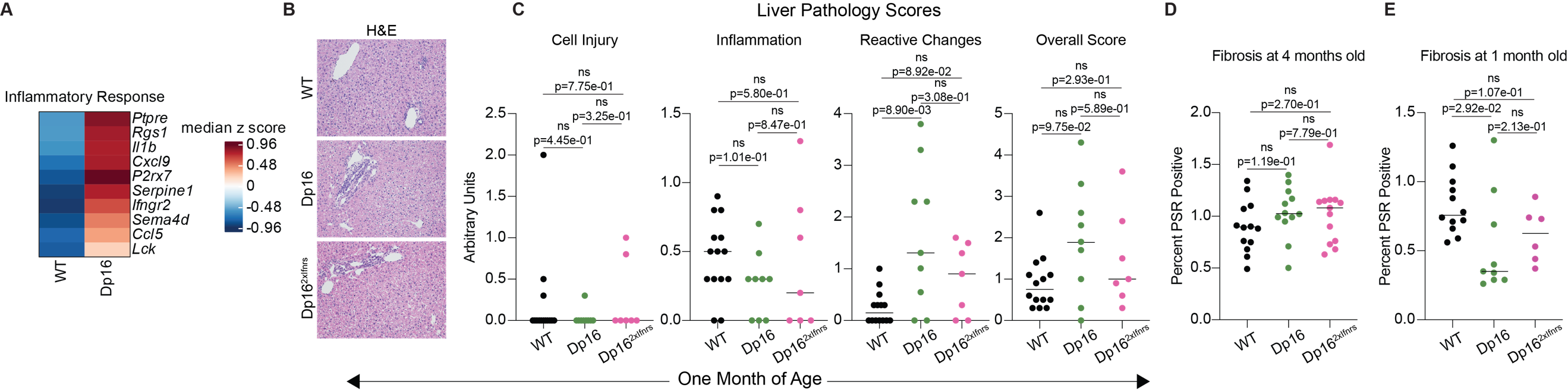

**Supplementary Figure 4. Elevated interferon signaling is not a major driver of Dp16 liver dysfunction.**

(**A**) Heatmap showing the median z score of the leading-edge genes from Inflammatory Response GSEA pathway in whole liver tissue. Color coding indicates a positive (red) or a negative (blue) median z score.

(**B**) Representative images of H&E-stained liver sections from adult wildtype, Dp16, and Dp16^2xIFNRs^ mice.

(**C**) Sina plots showing the metrics of liver pathology scoring from one month old wildtype (n=14, 7 females), Dp16 (n=9, 2 females), and Dp16^2xIFNRs^ (n=7, 4 females) mice.

(**D**) Sina plot showing the percent PSR positive pixels in wildtype (n=13, 6 females), Dp16 (n=12, 5 females), and Dp16^2xIFNRs^ (n=13, 7 females) mice.

(**E**) Sina plot showing the percent PSR positive pixels in one month old wildtype (n=12, 6 females), Dp16 (n=9, 2 females), and Dp16^2xIFNRs^ (n=6, 4 females) mice.

Within Sina plots, individual data are presented with a bar at the median and p-values, as determined by Mann-Whitney U test, are shown.

***Dunn et al., Supplementary Figure 5***

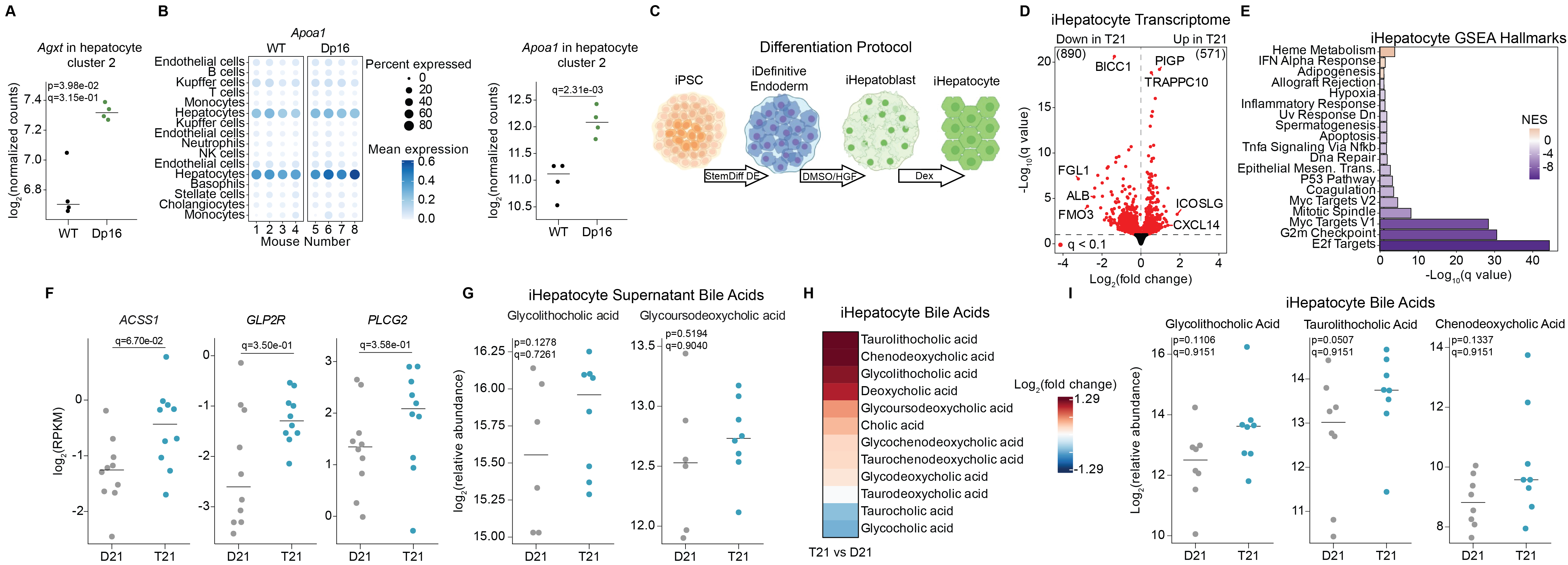

**Supplementary** **Figure 5. Liver dysfunction in Down syndrome is driven by hepatocyte dysfunction.**

(**A**) Sina plot displaying the log_2_ of the normalized counts of *Agxt* expression within hepatocyte cluster 2 in both wildtype and Dp16 animals. Individual data are presented with a bar at the median and p-values, as determined by Mann-Whitney U test, are shown.

(**B**) Bubble plot displaying *Apoa1* single-cell expression across clusters and animals, with color representing mean expression and size representing percent of cells with expression*.* Sina plot displays the log_2_ of the normalized pseudobulk counts of *Apoa1* expression within hepatocyte cluster 2 in both wildtype and Dp16 animals.

(**C**) Schematic illustrating the iPSC derived hepatocyte differentiation protocol.

(**D**) Volcano plot summarizing results of transcriptome analysis of iPSC derived hepatocytes (iHepatocytes). Significantly differentially expressed genes(q < 0.1) are colored in red.

(**E**) Barplot summarizing the results of Gene Set Enrichment Analysis of gene expression changes in iHepatocytes.

(**F**) Sina plots displaying the log_2_(RPKM) of various genes in D21 and T21 iHepatocytes.

(**G**) Sina plots displaying the log_2_(relative abundance) of various bile acids in the supernatant D21 and T21 iHepatocytes. Individual data are presented with a bar at the median and p-values, as determined by Mann-Whitney U test, are shown.

(**H**) Heatmap displaying the log_2_(fold change) of bile acids detected in trisomic versus disomic iHepatocytes. Color coding indicates a positive (red) or a negative (blue) log_2_(FoldChange) value. Asterisks denote significance (q<0.1).

(**I**) Sina plots displaying the log_2_(relative abundance) of various bile acids in D21 and T21 iHepatocytes. Individual data are presented with a bar at the median and p-values, as determined by Mann-Whitney U test, are shown.

***Dunn et al., Supplementary Figure 6***

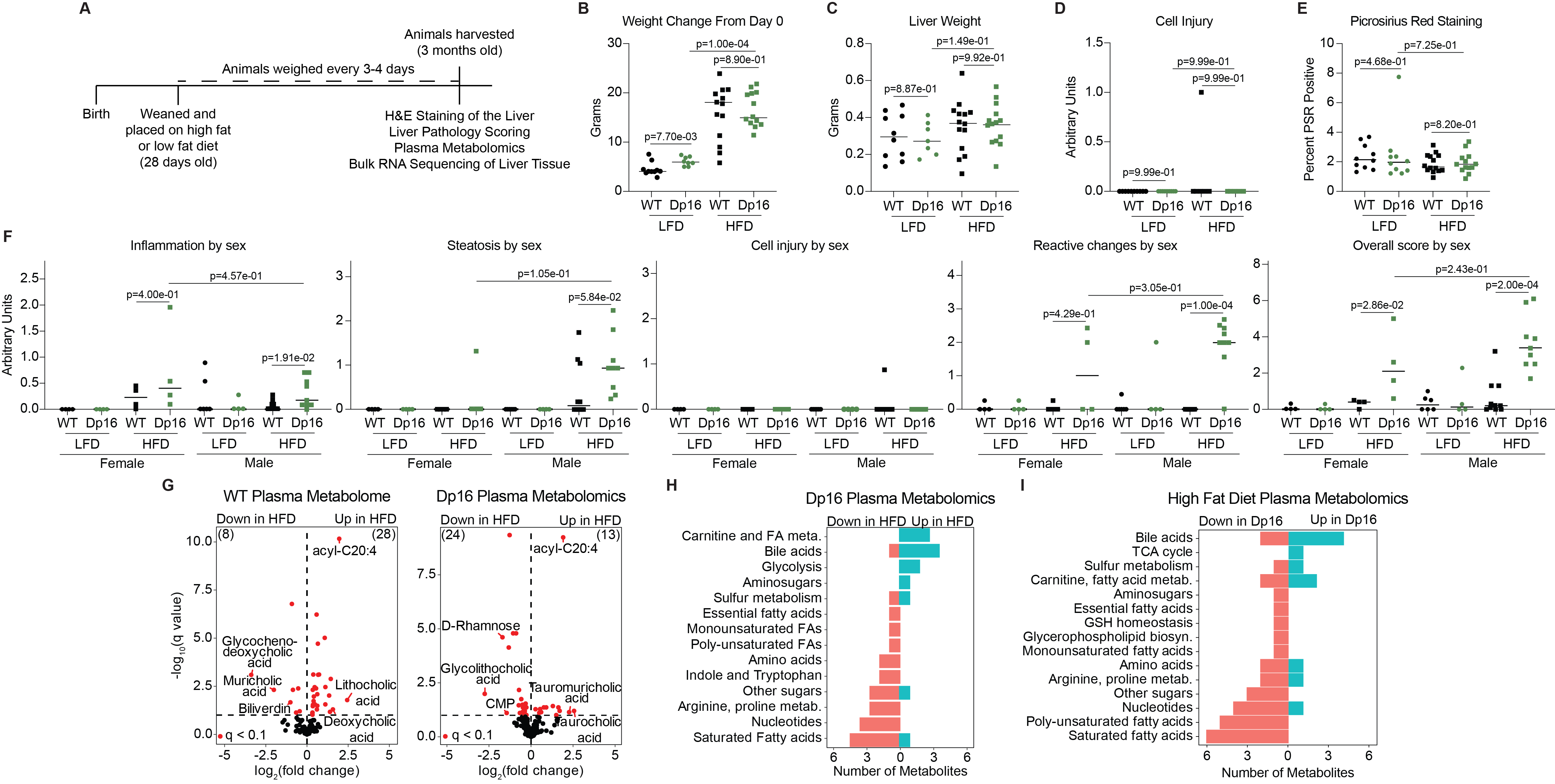

**Supplemental Figure 6. Liver pathology in Dp16 mice is dependent on diet.**

(**A**) Timeline showing high-fat diet experiment details.

(**B**) Sina plot displaying the amount of weight in grams animals gained over the course of treatment (wildtype LFD n=10, 4 females, Dp16 LFD n=8, 4 females, wildtype HFD n=14, 4 females, Dp16 HFD n=13, 4 females).

(**C**) Sina plot displaying the liver weight in grams for all treatment groups upon euthanasia.

(**D**) Sina plot displaying the cell injury liver pathology scoring parameter for all treatment groups.

(**E**) Sina plot showing the percent PSR positive pixels for all treatment groups.

(**F**) Sina plots displaying the liver pathology scores of LFD and HFD animals, broken out by sex.

(**G**) Volcano plots summarizing the results of plasma metabolomic analyses of WT LFD versus WT HFD mice and Dp16 HFD versus Dp16 LFD (wildtype LFD n=10, 4 females, Dp16 LFD n=8, 4 females, wildtype HFD n=13, 4 females, Dp16 HFD n=13, 4 females). Significant correlations (q < 0.1) are colored in red.

(**H**) Waterfall plot summarizing results of metabolite set enrichment analysis of plasma metabolites from Dp16 HFD versus Dp16 LFD mice. Bars are color-coded by increased relative abundance in DS (blue) versus decreased in DS (red).

(**I**) Waterfall plot summarizing results of metabolite set enrichment analysis of plasma metabolites from WT HFD versus Dp16 HFD mice. Bars are color-coded by increased relative abundance in DS (blue) versus decreased in DS (red).

Individual data are presented with a bar at the median and p-values, as determined by Mann-Whitney U test, are shown on all Sina plots.

***Dunn et al., Supplementary Figure 7***

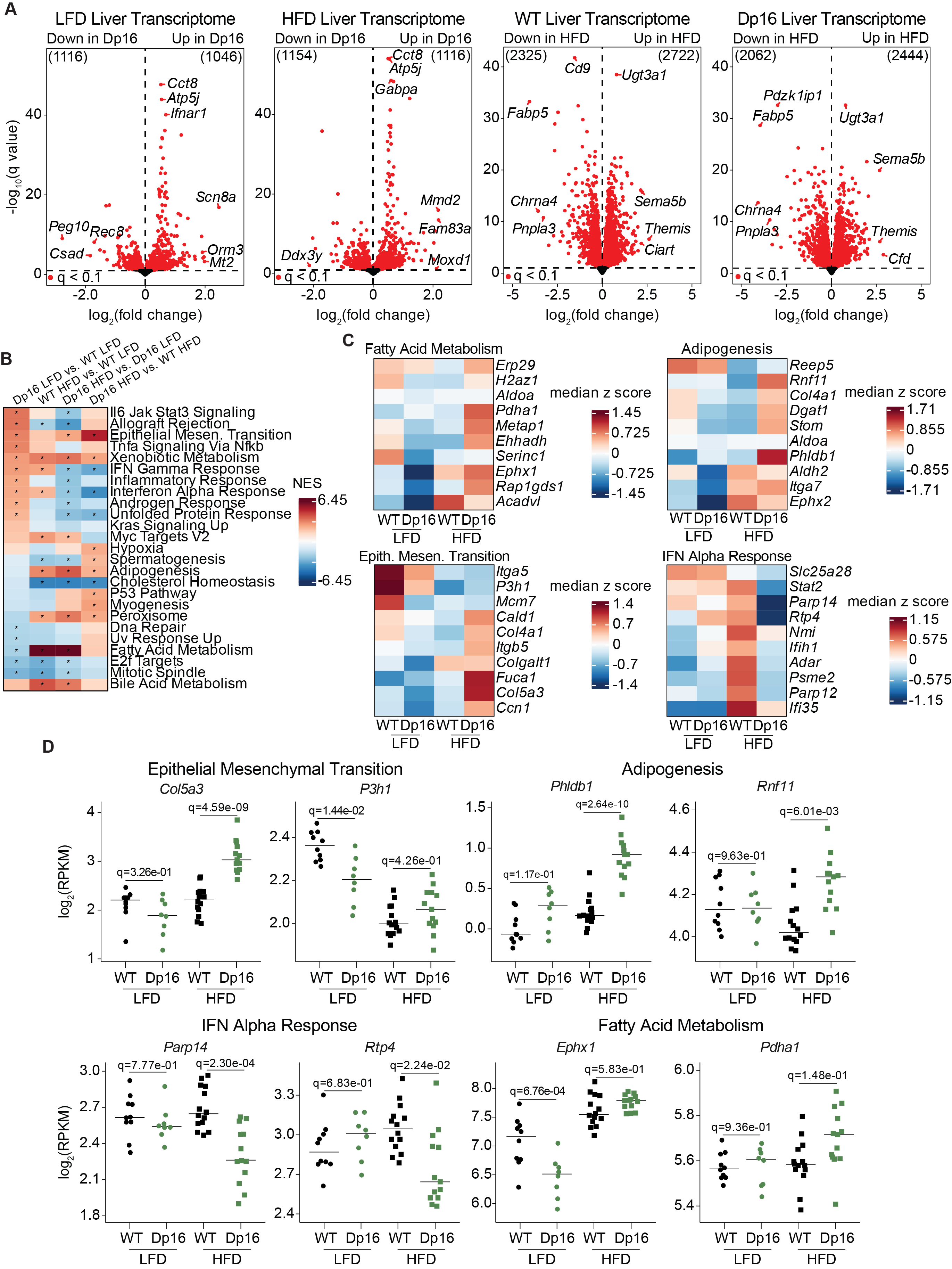

**Supplemental Figure 7.** **A high-fat diet contributes to global transcriptomic changes in the Dp16 liver.**

(**A**) Volcano plots summarizing results of whole liver transcriptome analysis of low-fat diet WT versus low-fat diet Dp16 mice, high-fat diet WT versus high-fat diet Dp16 mice, low-fat diet WT versus high-fat diet WT mice, and low-fat diet Dp16 versus high-fat diet Dp16 mice. Significantly differentially expressed (q<0.1) genes are colored in red.

(**B**) Heatmap displaying the enrichment scores of various Gene Set Enrichment Analysis signatures in four different comparisons. Asterisks denote significance (q<0.1).

(**C**) Heatmaps displaying the median z score of four different comparisons of the top 10 leading-edge genes for various Gene Set Enrichment Analysis signatures. Color coding indicates a positive (red) or a negative (blue) median z score.

(**D**) Sina plots displaying the log_2_(RPKM) of various genes from various Gene Set Enrichment Analysis signatures. Individual data are presented with a bar at the median and p-values, as determined by Mann-Whitney U test, are shown.

**Supplemental File Legends**

**Supplementary File 1. Human and Mouse Plasma Metabolomics.**

**Tab A.** Results of human plasma metabolomic analyses comparing those with Trisomy 21 (n=316) and controls (n=103).

Column description: Column A is the metabolite, B is the fold change, C is the log_2_(Fold Change), D is the unadjusted p-value, E is the Bonferroni-Hochberg adjusted p-value, and F is the linear regression model used.

**Tab B.** Results of correlation analysis of the relative abundance of taurolithocholic acid and BMI, as well as age.

Column description: column A is the internal participant ID, B is the age in years, C is the karyotype, D is the sex, batch is the sample batch, F is the weight in kilograms, G is the height in centimeters, H is the body mass index, I is the age, sex, and batch adjusted relative abundance of taurolithocholic acid, and J is the the BMI, sex, and batch adjusted relative abundance of taurolithocholic acid.

**Tab C.** Results of mouse plasma metabolomic analyses comparing Dp16 mice (n=13) to WT (n=9) mice.

Column description: Column A is the metabolite, B is the compound ID, C is the fold change, D is the log_2_(Fold Change), E is the unadjusted p-value, F is the Bonferroni-Hochberg adjusted p-value, and G is the linear regression model used.

**Supplementary File 2. Human and Proteomics and Mouse Liver Pathology Scoring.**

**Tab A.** Results of SOMAscan proteomic analysis of human plasma from individuals with DS (n=316) compared to controls (n=103).

Column description: Column A is the aptamer, B is the gene symbol, C is the fold change, D is the is the log_2_(Fold Change), E is the unadjusted p-value, F is the Bonferroni-Hochberg adjusted p-value, and G is the linear regression model used.

**Tab B.** Results of GSEA of SOMAscan proteomic data from human plasma from individuals with DS compared to controls.

Column description: Column A is the pathway, B is the unadjusted p-value, C is the Bonferroni-Hochberg adjusted p-value, D is the enrichment score, E is the normalized enrichment score, F is the size of the signature, and G is the leading-edge list.

**Tab C.** Results of GSEA Fibrosis and Steatosis signatures of SOMAscan proteomic data from human plasma from individuals with DS compared to controls.

Column description: Column A is the pathway, B is the unadjusted p-value, C is the Bonferroni-Hochberg adjusted p-value, D is the enrichment score, E is the normalized enrichment score, F is the size of the signature, and G is the leading-edge list.

**Tab D.** Results of WT and Dp16 mouse plasma ALT and AST quantification

Column description: Column A is the animal ID, B is the ALT concentration in units per liter, C is the AST concentration in units per liter, and D is the animal genotype and sex.

**Tab E.** Results of WT and Dp16 mouse plasma ALB quantification

Column description: Column A is the animal ID, B is the genotype, C is the sex, D is the age at harvest in days, E is the age at harvest in weeks, F is the litter the animal belonged to, and G is the albumin concentration in grams per deciliter.

**Tab F.** Results of WT and Dp16 mouse plasma IGF1 quantification

Column description: Column A is the animal ID, B is the genotype, C is the sex, D is the age at harvest in days, E is the age at harvest in weeks, and F is the IGF1 abundance.

**Tab G.** Results of WT and Dp16 mouse plasma CHI3L1 quantification

Column description: Column A is the animal ID, B is the genotype, C is the sex, D is the age at harvest in days, E is the age at harvest in weeks, and F is the litter the animal belonged to, and G is the CHI3L1 abundance.

**Tab H.** Results of WT and Dp16 mouse liver pathology scoring and Picrosirius Red (PSR) staining quantification

Column description: Column A is the animal ID, B is the genotype, C is the sex, D is the age at harvest in days, E is the age at harvest in weeks, F is the litter the animal belonged to, G is the total score of liver pathology scoring, H is the steatosis score, I is the inflammation score, J is the reactive changes score, K is the cell injury score, and L is the PSR quantification as a metric of total positive PSR.

**Tab I.** Results of 1 month old WT and Dp16 mouse liver pathology scoring and Picrosirius Red (PSR) staining quantification

Column description: Column A is the animal ID, B is the genotype, C is the sex, D is the age at harvest in days, E is the age at harvest in weeks, F is the litter the animal belonged to, G is the total score of liver pathology scoring, H is the steatosis score, I is the inflammation score, J is the reactive changes score, K is the cell injury score, and L is the PSR quantification as a metric of total positive PSR.

**Tab J.** Results of WT and Dp16 mouse plasma SAA quantification

Column description: Column A is the animal ID, B is the genotype, C is the sex, D is the age at harvest in days, E is the age at harvest in weeks, F is the litter the animal belonged to, and G is the SAA abundance.

**Tab K.** Results of 1 month old WT and Dp16 mouse plasma ALP quantification

Column description: Column A is the animal ID, B is the genotype, C is the sex, D is the age at harvest in days, E is the age at harvest in weeks, F is the litter the animal belonged to, and G is the ALP abundance in units per liter.

**Tab L.** Results of 1 month old WT and Dp16 mouse plasma CREA quantification

Column description: Column A is the animal ID, B is the genotype, C is the sex, D is the age at harvest in days, E is the age at harvest in weeks, F is the litter the animal belonged to, and G is the CREA abundance.

**Tab M.** Results of 1 month old WT and Dp16 mouse plasma SAA quantification

Column description: Column A is the animal ID, B is the genotype, C is the sex, D is the age at harvest in days, E is the age at harvest in weeks, F is the litter the animal belonged to, and G is the SAA abundance.

**Tab N.** Results of 1 month old WT and Dp16 mouse plasma CHI3L1 quantification

Column description: Column A is the animal ID, B is the genotype, C is the sex, D is the age at harvest in days, E is the age at harvest in weeks, F is the litter the animal belonged to, and G is the CHI3L1 abundance.

**Tab O.** Results of 1 month old WT and Dp16 mouse plasma CRP quantification

Column description: Column A is the animal ID, B is the genotype, C is the sex, D is the age at harvest in days, E is the age at harvest in weeks, F is the litter the animal belonged to, and G is the CRP abundance.

**Tab P.** Results of 1 month old WT and Dp16 mouse plasma ALB quantification

Column description: Column A is the animal ID, B is the genotype, C is the sex, D is the age at harvest in days, E is the age at harvest in weeks, F is the litter the animal belonged to, and G is the ALB abundance.

**Supplementary File 3. Mouse Liver bulk RNA-sequencing data.**

**Tab A.** Results of DESeq2 differential expression analysis of bulk RNA-sequencing of the liver from 4 month-old Dp16 (n=6) and WT (n=6) mice.

Column description: column A is the mouse gene name, B is the mouse chromosome encoding each gene, C is the mouse Ensembl gene ID, D is the base mean of a genotype, column E is the fold change between genotypes, F is the log2 fold change between genotypes, G is the adjusted fold change value between genotypes, H is the log2 adjusted fold change value between genotypes, I is the p-value, J is the q-value using the Benjamini-Hochberg method, K is the starting location of the gene, L is the ending location, M is the gene type, and N is the MGI ID.

**Tab B.** GSEA of the mouse liver RNA-sequencing data

Column description: Column A is the Gene Set, B is the unadjusted p-value, C is the Bonferroni-Hochberg adjusted p-value, D is the enrichment score, E is the normalized enrichment score, F is the size of the signature, and G is the leading-edge list.

**Supplementary File 4. Mouse Liver single-cell RNA-sequencing data and liver multiplexed immunofluorescence staining quantification.**

**Tab A.** GSEA of the mouse liver single-cell RNA-sequencing data

Column description: Column A is the cluster, B is the Gene Set, C is the unadjusted p-value, D is the Bonferroni-Hochberg adjusted p-value, E is the enrichment score, F is the normalized enrichment score, G is the size of the signature, and H is the leading-edge list.

**Tabs B-Q.** Results of DESeq2 analysis using gene-level pseudobulk count data to identify differentially expressed genes within each cluster

Column description: Column A is the cluster, B is the GeneID, C is the base mean of a genotype, D is the statistical model used, E is the fold change between genotypes, F is the log2 fold change between genotypes, G is the adjusted fold change value between genotypes, H is the log2 adjusted fold change value between genotypes, I is the p-value, and J is the q-value using the Benjamini-Hochberg method.

**Tab R.** Results of cluster abundance analysis of the mouse liver single-cell RNA-sequencing data

Column description: Column A is the cluster, B is the fold change between genotypes, C is the unadjusted p-value, D is the adjusted fold change value between genotypes using the Benjamini–Hochberg (BH) method, and E is the beta regression model used.

**Tab S.** Mouse liver multiplexed immunofluorescence staining quantification

Column description: Column A is the number of fields imaged per animal, B is the slide ID, C is the genotype and sex, D is the percent of total cells that were double positive for SMA and Desmin, E is the percent of total cells that were positive for PDPN, F is the percent of total cells that were positive for Lyve1, G is the percent of total cells that were positive for Albumin, H is the percent of total cells that were double positive for Lyve1 and PDPN and negative for CK19, I is the percent of total cells that were positive for Lyve1 and negative for PDPN and CK19, and J is the percent of total cells that were positive for Desmin.

**Supplementary File 5. Liver pathology scoring and plasma metabolomics from the Dp16^2xIfnrs^ mouse model.**

**Tab A.** Results from liver pathology scoring of adult WT, Dp16, and Dp16^2xIfnrs^ mice

Column description: Column A is the animal ID, B is the genotype, C is the sex, D is the age at harvest in days, E is the age at harvest in weeks, F is the litter the animal belonged to, G is the total score of liver pathology scoring, H is the steatosis score, I is the inflammation score, J is the reactive changes score, K is the cell injury score, and L is the PSR quantification as a metric of total positive PSR.

**Tab B.** Results from liver pathology scoring of 1 month old WT, Dp16, and Dp16^2xIfnrs^ mice

Column description: Column A is the animal ID, B is the genotype, C is the sex, D is the age at harvest in days, E is the age at harvest in weeks, F is the total score of liver pathology scoring, G is the steatosis score, H is the inflammation score, I is the reactive changes score, and J is the cell injury score.

**Tab C.** Results of adult WT, Dp16, Dp16^2xIfnrs^ mouse plasma ALB quantification

Column description: Column A is the animal ID, B is the genotype, C is the sex, D is the age at harvest in days, E is the age at harvest in weeks, F is the litter the animal belonged to, and G is the ALB concentration.

**Tab D.** Results of liver PSR quantification from 1 month old WT, Dp16, and Dp16^2xIfnrs^ mice

Column description: Column A is the animal ID, B is the genotype, C is the sex, D is the age at harvest in days, E is the age at harvest in weeks, and F is the PSR quantification as a metric of total positive PSR.

**Tab E.** Results of plasma metabolomic analysis comparing Dp16^2xIfnrs^ and Dp16 mice

Column description: Column A is the metabolite, B is the fold change, C is the log_2_(Fold Change), D is the unadjusted p-value, E is the Bonferroni-Hochberg adjusted p-value, F is the genotype comparison, and G is the linear regression model used.

**Tab F.** Results of plasma metabolomic analysis comparing Dp16^2xIfnrs^ and WT mice

Column description: Column A is the metabolite, B is the fold change, C is the log_2_(Fold Change), D is the unadjusted p-value, E is the Bonferroni-Hochberg adjusted p-value, F is the genotype comparison, and G is the linear regression model used.

**Supplementary File 6. iHepatocyte qPCR, transcriptome, and metabolomics data.**

**Tab A.** Gene expression of albumin over the stages of iHepatocyte differentiation

Column description: Column A is the iPSC cell line (where C indicates control samples and D indicates Trisomy 21), B is the replicate, C is the differentiation day, D is the cycle threshold (CT) of the 18s ribosomal RNA gene, E is the albumin expression, F is the delta CT (the difference between column E and D), G is the control delta CT (the average delta CT of the control samples), H is the delta CT control average (the difference between columns F and G), I is the fold gene expression, and J is the average fold change.

**Tab B.** Gene expression of Cyp3A4 over the stages of iHepatocyte differentiation

Column description: Column A is the iPSC cell line (where C indicates control samples and D indicates Trisomy 21), B is the replicate, C is the differentiation day, D is the cycle threshold (CT) of the 18s ribosomal RNA gene, E is the Cyp3A4 expression, F is the delta CT (the difference between column E and D), G is the control delta CT (the average delta CT of the control samples), H is the delta CT control average (the difference between columns F and G), I is the fold gene expression, and J is the average fold change.

**Tab C.** Transcriptome analysis of iHepatocytes

Column description: Column A is the gene name, B is the chromosome encoding each gene, C is the Ensembl gene ID, D is the base mean of a karyotype, E is the SVA model used, F is the fold change between karyotypes, G is the log2 fold change between karyotypes, H is the adjusted fold change value between karyotypes using the Benjamini–Hochberg (BH) method, I is the log2 adjusted fold change value between karyotypes using the BH method, J is the p-value, K is the q-value using the BH method, L is the starting location of the gene, M is the ending location, N is the gene type, and O is the HGNC ID.

**Tab D.** GSEA of the iHepatocyte RNA-sequencing data

Column description: Column A is the pathway, B is the unadjusted p-value, C is the Bonferroni-Hochberg adjusted p-value, D is the enrichment score, E is the normalized enrichment score, F is the size of the signature, and G is the leading-edge list.

**Tab E.** iHepatocyte supernatant metabolomic analysis

Column description: Column A is the metabolite, B is the compound ID, C is the pathway the metabolite belongs to, D is the analyte group (metabolites versus bile acids), E is the fold change, F is the log_2_(Fold Change), G is the unadjusted p-value, and H is the Bonferroni-Hochberg adjusted p-value.

**Tab F.** iHepatocyte cell metabolomic analysis

Column description: Column A is the metabolite, B is the compound ID, C is the pathway the metabolite belongs to, D is the analyte group (metabolites versus bile acids), E is the fold change, F is the log_2_(Fold Change), G is the unadjusted p-value, and H is the Bonferroni-Hochberg adjusted p-value.

**Supplementary File 7.** **Animal weight change, liver pathology, and liver weights from animals fed a high-fat or low-fat diet.**

**Tab A.** Weight change of WT and Dp16 mice fed either a high-fat or low-fat diet over the course of treatment

Column description: Column A is the sex of the animal, B is the animal ID, C is the cage ID, D is the treatment, E is the genotype, F-AC are the animal weights in grams on the indicated dates, and AD is the change in weight from day 1.

**Tab B.** Liver weight upon euthanasia of WT and Dp16 mice fed either a high-fat or low-fat diet

Column description: Column A is the ID of the animal, and B is the liver weight in grams.

**Tab C.** Liver pathology scoring of WT and Dp16 mice fed either a high-fat or low-fat diet

Column description: Column A the animal ID, B is the genotype, C is the treatment group, D is the sex, E is the age at harvest in days, F is the age at harvest in weeks, G is the total score of liver pathology scoring, H is the steatosis score, I is the inflammation score, J is the reactive changes score, K is the cell injury score, and L is the PSR quantification as a metric of total positive PSR.

**Supplementary Figure 8. Metabolomic analyses of plasma from WT and Dp16 mice fed either a high-fat or low-fat diet.**

**Tab A.** Metabolomic analysis of plasma from Dp16 mice fed a high-fat diet compared to a low-fat diet

**Tab B.** Metabolomic analysis of plasma from WT mice fed a high-fat diet compared to a low-fat diet

**Tab C.** Metabolomic analysis of plasma from Dp16 mice compared to WT mice fed a high-fat diet

**Tab D.** Metabolomic analysis of plasma from Dp16 mice compared to WT mice fed a low-fat diet

Column description for all tabs: Column A is the metabolite, B is the compound ID, C fold change, D is the log_2_(Fold Change), E is the unadjusted p-value, F is the Bonferroni-Hochberg adjusted p-value, G is the linear regression model used, and H is the comparison group.

**Supplementary Figure 9.** **Transcriptome analysis of the liver from WT and Dp16 mice fed a high-fat or low-fat diet.**

**Tab A.** Transcriptome analysis of the liver from WT mice fed a high-fat diet compared to a low-fat diet

**Tab B.** Transcriptome analysis of the liver from Dp16 mice fed a high-fat diet compared to a low-fat diet

**Tab C.** Transcriptome analysis of the liver from Dp16 mice compared to WT mice fed a low-fat diet

**Tab D.** Transcriptome analysis of the liver from Dp16 mice compared to WT mice fed a high-fat diet

Column description for all tabs: column A is the mouse gene name, B is the mouse chromosome encoding each gene, C is the mouse Ensembl gene ID, D is the base mean of a genotype, column E is the fold change between genotypes, F is the log2 fold change between genotypes, G is the adjusted fold change value between genotypes, H is the log2 adjusted fold change value between genotypes, I is the p-value, J is the q-value using the Benjamini-Hochberg method, K is the starting location of the gene, L is the ending location, M is the MGI ID, N is the SVA model used, and O is the comparison groups.
